# Dopaminergic modulation of the exploration/exploitation trade-off in human decision-making

**DOI:** 10.1101/706176

**Authors:** Karima Chakroun, David Mathar, Antonius Wiehler, Florian Ganzer, Jan Peters

## Abstract

A central issue in reinforcement learning and decision-making is whether to exploit knowledge of reward values, or to explore novel options. Although it is widely hypothesized that dopamine neurotransmission plays a key role in regulating this balance, causal evidence for a role of dopamine in human exploration is still lacking. Here, we use a combination of computational modeling, pharmacological intervention and functional magnetic resonance imaging (fMRI) to test for a causal effect of dopamine transmission on the exploration-exploitation trade-off in humans. 31 healthy male subjects performed a restless four-armed bandit task in a within-subjects design under three drug conditions: 150mg of the dopamine precursor L-dopa, 2mg of the D2 receptor antagonist haloperidol, and placebo. In all conditions, choice behavior was best explained by an extension of an established Bayesian learning model accounting for perseveration, uncertainty-based exploration and random exploration. Uncertainty-based exploration was attenuated under L-dopa compared to placebo and haloperidol. There was no evidence for a modulation of prediction error signaling or categorical effects of exploration/exploitation under L-dopa, whereas model-based fMRI revealed that L-dopa attenuated neural representations of overall uncertainty in insula and dorsal anterior cingulate cortex. Our results highlight the computational role of these regions in exploration and suggest that dopamine modulates exploration by modulating how this circuit tracks accumulating uncertainty during decision-making.

## Introduction

A central aspect of a broad spectrum of decision problems is the weighting of when to exploit, i.e. to choose a familiar option with a well-known reward value, and when to explore, i.e. to try an alternative option with an uncertain but potentially higher payoff. This decision dilemma is commonly known as the “exploration/exploitation trade-off” (Cohen, McClure, & Yu, 2007; Addicott et al. 2017). Striking a good balance between exploration and exploitation is essential for maximizing rewards and minimizing costs in the long term (Addicott et al., 2017). Too much exploitation prevents an agent from gathering new information in an uncertain environment, fosters inflexibility and habit formation. Too much exploration, on the other hand, may lead to inefficient and inconsistent decision-making, thereby reducing long-term payoffs (Beeler 2014; Addicott 2017). Despite the high relevance of the explore/exploit trade-off for optimal decision-making, research is only beginning to unravel the mechanisms through which animals and humans solve this dilemma. Several cognitive task have been developed to test explore/exploit behavior in both animals and humans. The most widely used paradigm is the multi-armed bandit task (Robbins, 1952; Gittins & Jones, 1974). It mirrors a casino’s slot-machine with multiple arms. Several implementations exist that differ according to the number of arms and their underlying reward structure. The restless bandit paradigm uses continuous, slowly drifting rewards for each bandit that encourage participants to strike a balance between exploiting the currently best option and exploring alternative bandits to keep track of their evolving rewards (Daw et al., 2006; Addicott et al., 2017). Other prominent paradigms include the (patch) foraging task (Cook et al., 2013; Addicott, Pearson, Kaiser, Platt, & McClernon, 2015; Constantino & Daw, 2015) that mirrors exploration and exploitation of food sources in a more naturalistic setting, and the horizon task (Wilson, Geana, White, Ludvig & Cohen, 2014) that examines exploration in series of discrete games. These paradigms offer different approaches to measure explore/exploit behavior and may be used to address different research questions. Computational models have served as an elegant tool for modeling behavior on these tasks, yielding insights into latent cognitive processes, and inter individual differences (Sutton & Barto, 1998; Daw et al., 2006, 2011; Gershman 2018).

Computational algorithms modeling exploration behavior during learning have at least two components: a learning rule and a choice rule. The learning rule describes how subjective value estimates of an option’s mean outcome are updated for each choice option based on experience, e.g. via the classical ‘Delta rule’ (Rescorla Wagner, 1972) from reinforcement learning theory (Sutton & Barto, 1998). Work on explore/exploit behavior has also to utilized a ‘Bayesian learner’ model that relies on a Kalman filter model that simultaneously tracks estimates of outcome mean and uncertainty (e.g. Daw et al., 2006; Speekenbrink & Konstantinidis, 2015), and uses an uncertainty-dependent delta rule for value updating. The choice rule then accounts for how learned values give rise to choices. Here, exploration can be due to at least two mechanisms. First, exploration could result from a probabilistic selection of sub-optimal options as in ε-greedy or softmax choice rules (Sutton & Barto, 1998), henceforth referred to as “random exploration” (Daw et al., 2006; Speekenbrink & Konstantinidis, 2015). In contrast, exploration could also be based on the degree of uncertainty associated with an option (Daw et al., 2006; Wilson et al., 2014), such that highly uncertain options have a higher probability to be strategically explored by an agent, henceforth referred to as “uncertainty-based exploration”. However, estimation of exploration/exploitation behavior might be partially confounded by perseveration (i.e. repeating previous choices irrespective of value or uncertainty), a factor that has not been incorporated into previous models of explore/exploit behavior (Badre et al., 2012; Speekenbrink & Konstantinidis, 2015).

Dopamine (DA) neurotransmission is thought to play a central role in the explore/exploit trade-off. Striatal phasic DA release is tightly linked to reward learning based on reward prediction errors (RPE) (Steinberg, Keiflin, Boivin et al., 2013) that reflect differences between experienced and expected outcomes, and serve as a ‘teaching signal’ that update value predictions (Schultz, Dayan & Montague, 1999; Schultz, 2016; Tsai et al., 2009; Glimcher et al., 2011; Chang et al., 2018). Exploitation has been linked to polymorphisms in genes controlling striatal DA signaling, namely the DRD2 gene (Frank et al., 2009) predictive of striatal D2 receptor availability (Hirvonen, 2004), and the DARPP-32 gene involved in striatal D1 receptor-mediated synaptic plasticity and reward learning (e.g. Calabresi, 2000; Stipanovich, 2008). Variation in the slower tonic DA signal might also contribute to an adaptive regulation of exploration/exploitation. Beeler et al. (2010) found that dopamine-transporter (DAT) knockdown mice that are characterized by increased striatal levels of tonic DA (Zhuang et. al, 2001) showed higher random exploration compared to wild-type controls. In addition to striatal DA, prefrontal DA might also be involved in explore/exploit behavior. In humans, both uncertainty-based and random exploration have been associated with variations in the catechol-O-methyltransferase (COMT) gene (Kayser et al., 2015; Gershman & Tzovaras, 2018) that modulates prefrontal DAergic tone (Meyer-Lindenberg 2005). Participants with putatively higher prefrontal DA tone had highest levels of exploration. Within a model-based/model-free reinforcement learning framework (Dolan & Dayan, 2013), exploitation can be regarded as a model-free learning strategy, whereas uncertainty-based exploration arguably constitutes a model-based strategy as it relies on an internal model of the environment (the Gaussian random walks that constitute the tasks payoff structure). Model-based learning is related to striatal presynaptic DA levels as assessed by [(18)F]DOPA positron emission tomography (Deserno et al., 2015).

These findings regarding the roles of striatal and frontal DA in exploration resonate with cognitive neuroscience studies suggesting that exploration and exploitation rely on distinct neural systems. Daw et al. (2006) showed that frontopolar cortex (FPC) is activated during exploratory choices, possibly facilitating behavioral switching between an exploitative and exploratory mode by overriding value-driven choice tendencies (Daw et al., 2006; Badre et al., 2012; Addicott et al., 2014; Mansouri et al., 2017). In line with this idea, up- and down-regulation of FPC excitability via transcranial direct current stimulation (TDCS) increases and decreases exploration during reward-based learning (Becharelle et al., 2015). Anterior cingulate cortex (ACC) and anterior insula (AI), have also been implicated in exploration (Addicott et al., 2014; Laureiro-Martínez et al., 2014, 2015; Blanchard & Gershman, 2018), although their precise computational role remains elusive (Blanchard & Gershman, 2018). Both regions may trigger attentional reallocation to salient choice options in the light of increasing uncertainty (Laureiro-Martínez, 2015). Exploitation, on the other hand, is thought to be predominantly supported by structures within a “valuation” network including ventromedial prefrontal cortex (vmPFC), orbitofrontal cortex (OFC), ventral striatum and hippocampus (Daw et al., 2006; Batra & Kable, 2013; Clithero & Rangel, 2014; Laureiro-Martínez et al., 2014, 2015).

Despite these advances, evidence for a causal link between DA transmission, exploration and the underlying neural mechanisms in humans is still lacking. To address this issue, we combined computational modeling and functional magnetic resonance imaging (fMRI) with a pharmacological intervention in a double-blind, counterbalanced, placebo-controlled within-subjects study. Participants performed a restless four-armed bandit task (Daw et al., 2006) during fMRI under three drug conditions: the DA precursor L-dopa (150mg), the DRD2 antagonist haloperidol (2mg), and placebo. While L-dopa is thought to stimulate DA transmission by providing increased substrate for DA synthesis, haloperidol reduces DA transmission by blocking D2 receptors. We extended previous modeling approaches of exploration behavior (Daw et al., 2006; Speekenbrink & Konstantinidis, 2015) using a hierarchical Bayesian estimation scheme. Specifically, we jointly examined dopaminergic drug-effects on uncertainty-based and random exploration as well as perseveration, hypothesizing that choice behavior would be best accounted for by a model that accounts for all three processes (Schönberg, Daw, Joel, & O’Doherty, 2007; Rutledge et al., 2009; Payzan-LeNestour & Bossaerts, 2012). We hypothesized both random and uncertainty-based exploration to increase under L-dopa and decrease under haloperidol compared to placebo (Frank et al., 2009; Beele et al., 2010; Gershman & Tzovaras, 2018). We further hypothesized that this would be accompanied by a corresponding modulation of brain activity in regions implicated in exploration, foremost the FPC, ACC and AI (Daw et al., 2006; Becharelle et al., 2015; Blanchard & Gershman, 2018). To examine potential individual differences in dopaminergic drug effects we also collected putative proxy measures of DA function, namely spontaneous eye blink-rate (sEBR) (Jongkees and Colzato, 2016) and working memory capacity (WMC) (Cools et al., 2008; Cools & D’Esposito, 2011) in a separate ‘baseline-session’. We predicted that explore/exploit behavior and their drug-related changes would be modulated by the individual DA baseline, as indexed by our DA proxy measures, according to an inverted-u-shaped function (Cools & D’Esposito, 2011; Cavanagh et al., 2014).

## Results

### Choice behavior contains signatures of uncertainty-based exploration and perseveration

On each testing day, separated by exactly one week, participants performed 300 trials of a 4-armed restless bandit task (Daw et al., 2006, see Methods section) during fMRI, under three pharmacological conditions (Placebo, Haloperidol, L-DOPA). We first examined whether choice behavior in the three sessions indeed contained signatures of random exploration, uncertainty-based exploration and perseveration. To this end, we set up six separate computational models that differed regarding the implemented learning and choice rules within a hierarchical Bayesian framework using the STAN modeling language (version 2.17.0; Stan Development Team, 2017). We compared two learning rules: the classical Delta rule from temporal-difference algorithms (e.g. Sutton & Barto, 1998), and a Bayesian learner (Daw et al. (2006)) that formalizes the updating process with a Kalman filter (Kalman et al., 1960). In the former model, values are updated based on prediction errors that are weighted with a constant learning rate. In contrast, the Kalman filter additionally tracks the uncertainty of each bandit’s value, and value updating is proportional to the uncertainty of the chosen bandit (Kalman gain, see Methods section). These learning rules were combined with three different choice rules that were all based on a softmax action selection rule (Sutton & Barto, 1998; Daw et al., 2006). Choice rule 1 was a standard softmax with a single inverse temperature parameter (β) modeling random exploration. Choice rule 2 included an additional free parameter (*φ*) modeling an exploration bonus that scaled with the estimated uncertainty of the chosen bandit (uncertainty-based exploration). Finally, the third choice rule included an additional free parameter (ρ) modeling a perseveration bonus for the bandit chosen on the previous trial. Leave-one-out (LOO) cross-validation estimates (Vehtari, Gelman, & Gabry, 2017) were computed over all drug-conditions, and for each condition separately to assess the models’ predictive accuracies. The Bayesian learning model with terms for uncertainty-based exploration and perseveration showed highest predictive accuracy in each drug condition and overall (Figure 1).

**Figure 1.**
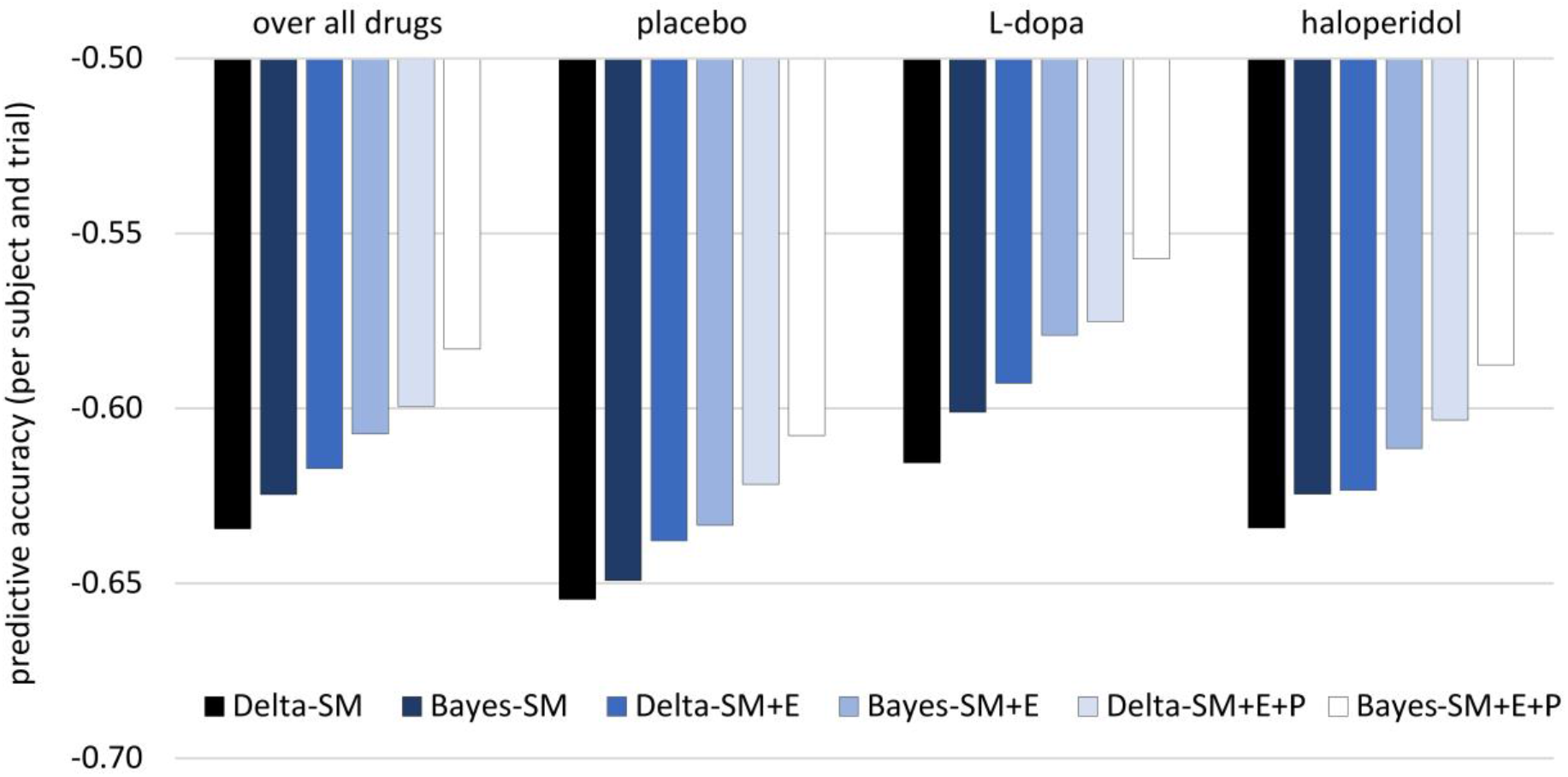
Results of the cognitive model comparison. Leave-one-out (LOO) estimates were calculated over all drug conditions (n=31 subjects with t=3 *300 trials) and once separately for each drug condition (n=31 with t=300). All LOO estimates were devided by the total number of data points in the sample (n*t) for better comparability across the different approaches. Note that the relative order of LOO estimates is invariant to linear transformations. Delta: simple delta learning rule; Bayes: Bayesian learner; SM: softmax (random exploration); E: uncertainty-based exploration; P: perseveration.

### Accounting for perseveration boosts estimates of uncertainty-based exploration

If perseveration is not explicitly accounted for, variance attributable to perseveration might influence the uncertainty-based exploration parameter (*φ*) in the form of an uncertainty-avoiding choice bias (Badre et al., 2012; Payzan-LeNestour & Bossaerts, 2012). Following up on this, we compared φ estimates for the placebo condition between the winning Bayesian learning model and the reduced model without a perseveration term. Subject-level medians for *φ* were highly correlated between models (r_29_=.90, p<.001), but were significantly higher for the model including a perseveration term than for the reduced model (mean difference=0.79, paired t-test: t_30_=7.97, p < .001). This was also true for the corresponding group-level mean parameter of *φ* (full model: 0.95; reduced model: 0.16). The number of subjects who showed a negative median *φ,* reflecting a discouragement rather than an encouragement of uncertainty-based exploration was also reduced (full model 6/31; reduced model 13/31). Together, this shows that explicitly accounting for perseveration improved sensitivity to detect effects of uncertainty-based exploration.

### Model-based regressors and classification of exploration trials

In the best-fitting Bayesian model, participants’ choices are stochastically dependent on three factors: the prior belief of the mean reward value of each bandit (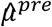; Figure 2a), the exploration bonus, i.e. the prior belief of each bandit’s payout variance (‘uncertainty’) scaled with the exploration bonus parameter *φ* (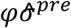; Figure 2b), and the perseveration bonus (*I*ρ; where *I* denotes an indicator function with respect to the bandit chosen on the previous trial; Figure 2c). Based on these quantities, which are computed for each bandit, the model computes the choice probabilities for all four bandits on each trial (*P*; Figure 2d, See Eq. (7) in the Methods section). Between trials, participants’ prior belief of the chosen bandit’s mean reward value is updated according to the reward prediction error (*δ*, Figure 2e), as the difference between their prior belief and the actual reward outcome of the chosen bandit. Based on the model, participants’ choices can be classified as exploitation (i.e. when the bandit with highest expected value was selected), or exploration (i.e. when any other bandit was selected) (Daw et al., 2006). We extended this binary classification of Daw et al. (2006) by further dividing exploration trials into uncertainty-based exploration (i.e., trials where the bandit with the highest exploration bonus was chosen), and random exploration trials (i.e. trials where one of the remaining bandits was chosen). The trinary classification scheme corresponded well with the respective model parameters (β, *φ,* ρ; see *SI*) Over trials, the summed uncertainty over all bandits (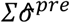; Figure 2f) fluctuates in relation to the fraction of exploration.

**Figure 2.**
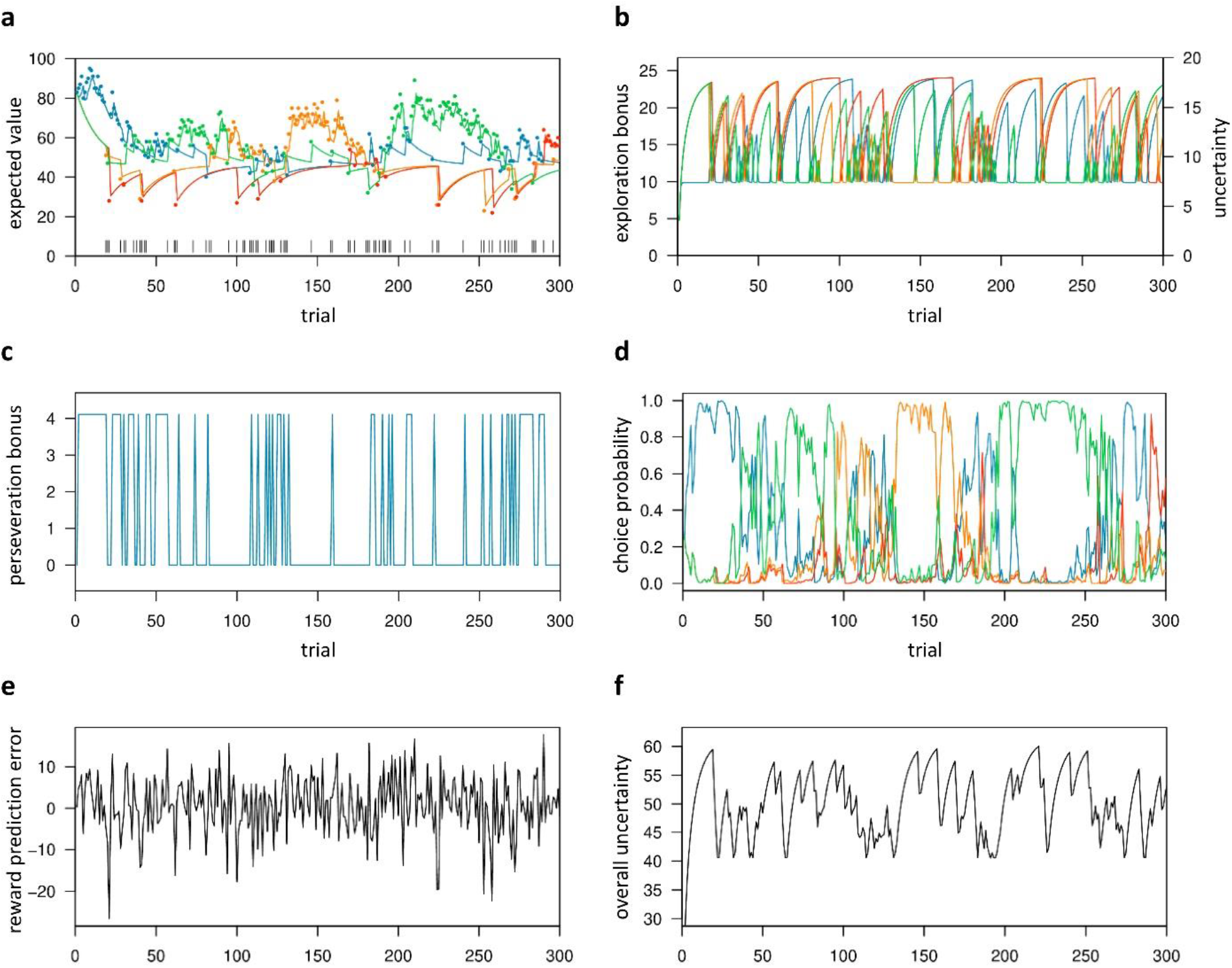
Trial-by-trial variables of the best-fitting Bayesian learning including both random and uncertainty-based exploration and perseveration. Trial-by-trial estimates are shown for the placebo data of one representative subject with posterior medians: β=0.29, *φ* =1.34, and ρ=4.11. (a) Colored lines depict the expected values 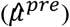 of the four bandits, whereas colored dots denote actual payoffs. Vertical black lines mark trials classified as exploratory (Daw et al., 2006). (b) Exploration bonus 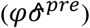 and uncertainty 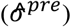 for each bandit. (c) Perseveration bonus (*I*ρ). This bonus is a fixed value added only to the bandit chosen in the previous trial, shown here for one bandit. (d) Choice probability (*P*). Each colored line represents one bandit. (e) Reward prediction error (*δ*). (f) The subject’s overall uncertainty 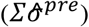, i.e. the summed uncertainty over all four bandits.

### L-dopa reduces uncertainty-based exploration

Dopaminergic drug effects were first examined for the group-level (mean *M* posteriors for β (random exploration), *φ* (uncertainty-based exploration) and ρ (perseveration)) of the best-fitting full Bayesian model. Posterior distributions for these parameters were estimated separately for each drug condition (Figure 3a). Under L-dopa, the posterior group-level mean of *φ* (*M^φ^*) was substantially reduced compared to both placebo and haloperidol (Figure 3a), such that the 90% highest density intervals of the difference of the posterior distributions of *M^φ^* (Figure 3b) did not overlap with zero (see *SI* for details).

**Figure 3.**
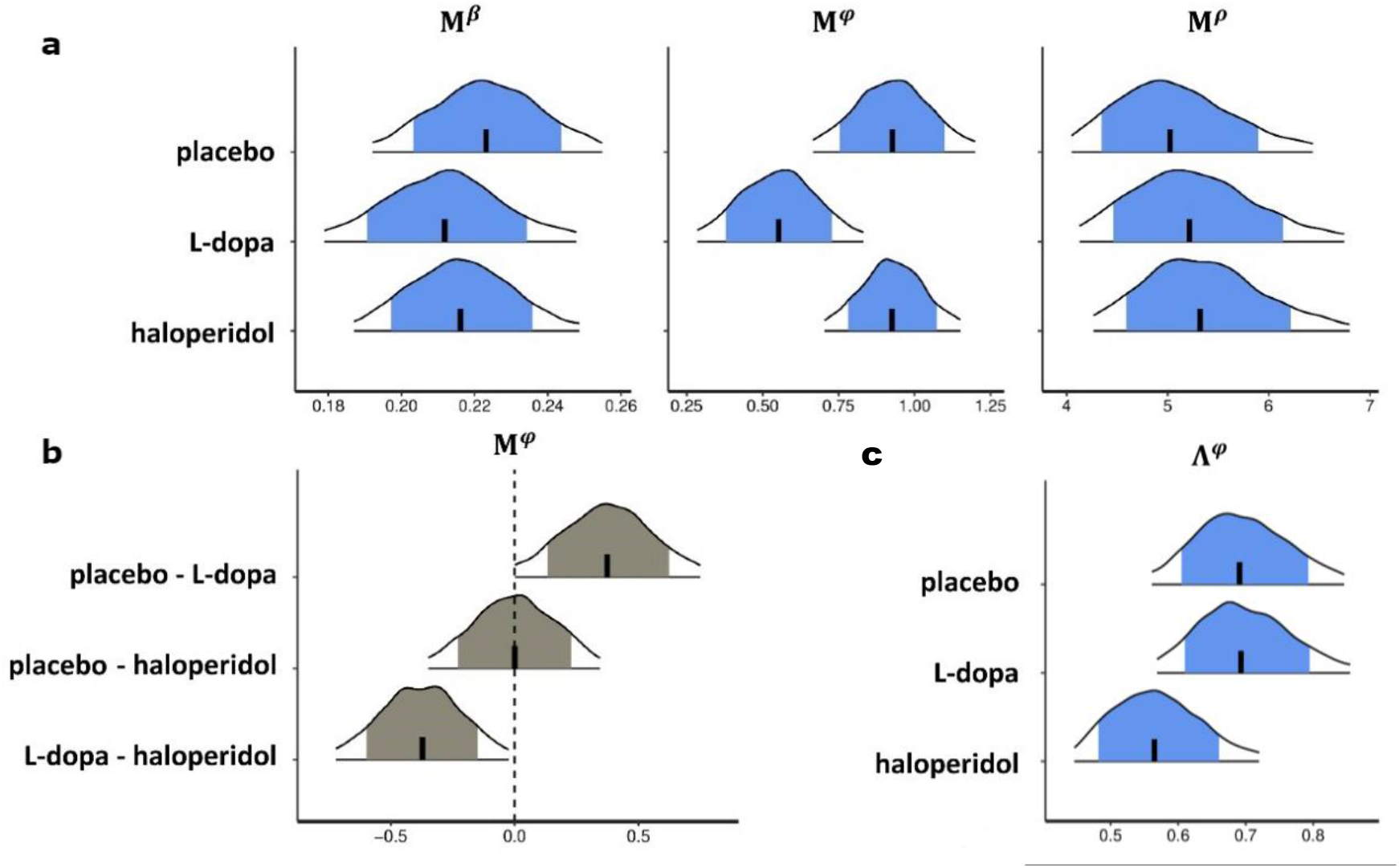
Drug effects for the group-level parameter estimates of the best-fitting Bayesian model. (a) Shown are posterior distributions of the group-level mean (*M*) of all choice parameters (β, *φ*, ρ), separately for each drug condition, and (b) posterior drug differences of the group-level mean (M) of the parameter *φ,* as well as (c) posterior distributions of the group-level standard deviation (Λ) of parameter *φ*. Each plot shows the median (vertical black line), the 80 % central interval (blue (grey) area), and the 95 % central interval (black contours); β: random exploration, *φ*: uncertainty-based exploration; ρ: perseveration parameter. For drug effects on the standard deviation of the group-level median parameters *β* and ρ see *SI.*

In contrast, we did not observe effects of L-dopa on random exploration (β, Figure 3a) or perseveration (ρ, Figure 3a). Somewhat surprisingly, haloperidol showed no effects on the posterior group-level means of model parameters. Instead, haloperidol appeared to decrease the standard deviation (*Λ*) of the posterior group-level means of *φ* to some extent (placebo: 0.69, L-dopa: 0.70, haloperidol: 0.57); Figure 3c). For illustrative purposes, we next examined drug effects on the subject-level parameters (β, *φ,* ρ). In the L-dopa condition individual parameters for *φ* were reduced compared to placebo and haloperidol (Figure 4). We did not observe such an effect under haloperidol. However, in line with the attenuated standard deviation of the group-level *φ* medians, haloperidol reduced the variability of the subject-level *φ* medians (range=[-0.43, 2.33]; SD=0.64) compared to placebo (range=[-0.95, 2.48]; SD=0.85) and L-dopa (range=[-1.77, 2.00]; SD=0.85). Visual inspection suggested that haloperidol increased *φ* for subjects with a relatively low value under placebo and decreased *φ* for subjects with a relatively high *φ* value under placebo (Figure 4). We observed similar results in an additional analysis of drug-effects on the percentage of exploitation and exploration trials (overall, random and uncertainty-based) per subject (see *SI*). We found no evidence for drug effects on model-free measures of choice behavior (see *SI*).

**Figure 4.**
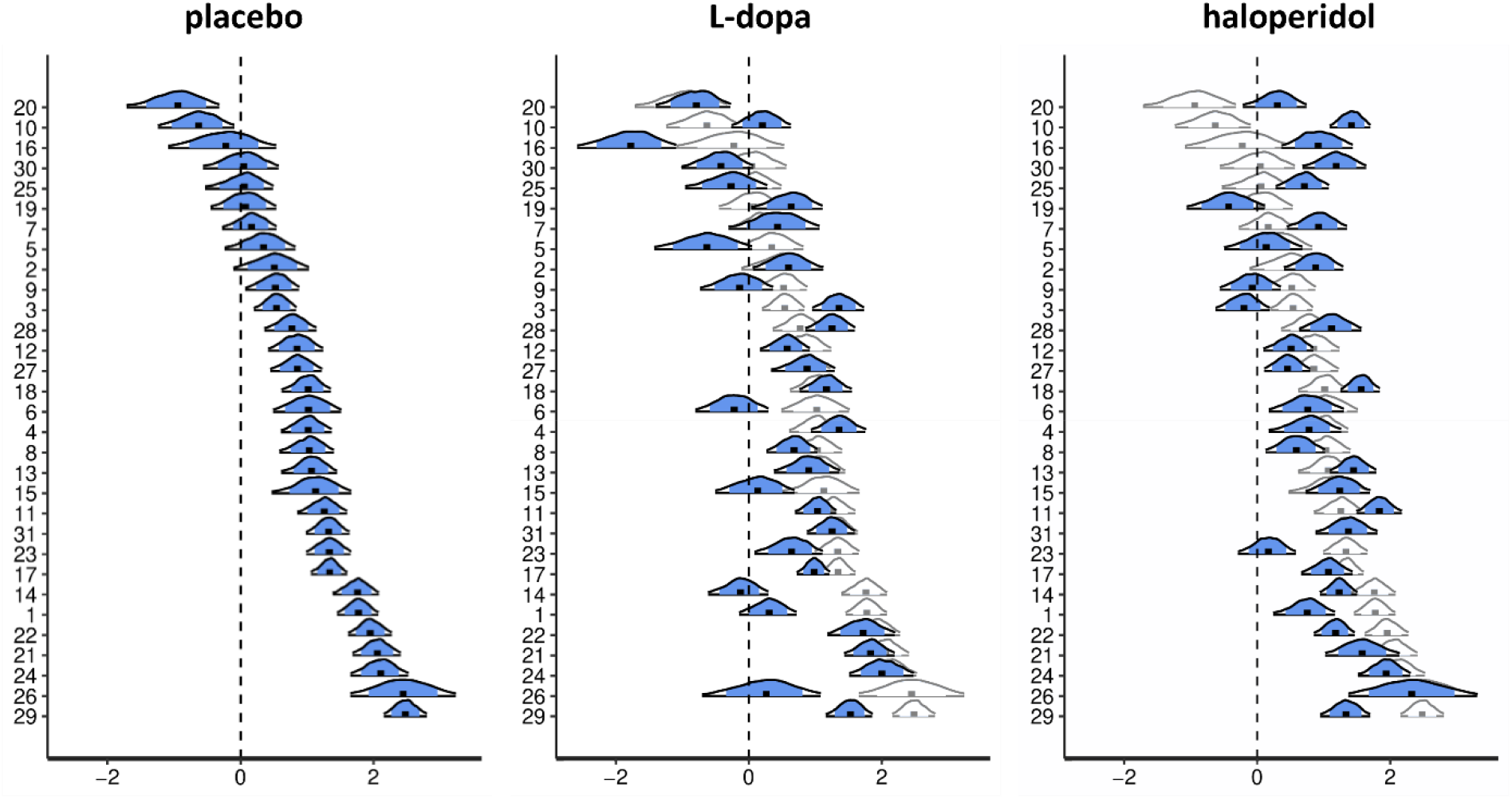
Drug effects for the subject-level parameter estimates of the uncertainty-based exploration parameter *φ*. Shown are posterior distributions of the subject-level parameter *φ* from the best-fitting Bayesian model, separately for each drug condition. Each plot shows the median (black dot), the 80 % central interval (blue area), and the 95 % central interval (black contours). For the L-dopa and haloperidol conditions, posterior distributions (in blue) are overlaid on the posterior distributions of the placebo condition (in white) for better comparison.

### No evidence for the inverted-U-shaped dopamine hypothesis

According to the inverted-U-shaped DA hypotheses (Cools & D’Esposito (2011), we predicted participants’ individual DA baseline to modulate explore/exploit behavior in a quadratic fashion and to linearly predict the strength and direction of drug-related effects. As potential proxies for subjects’ DA baseline, we assessed spontaneous blink rate (Jongkees and Colzato, 2016) and working memory capacity via three working memory tasks (Operation/Rotation/Listening SPAN) (Cools & D’Esposito, 2011; Kane et al., 2004; Unsworth, Redick, Heitz, Broadway, & Engle, 2009; Redick et al., 2012). Scores on the first component of a principal component analysis (PCA) (explaining 56.6% of the shared variance) over the z-transformed working memory task scores were used as a working memory compound score. We then used regression to test for linear and quadratic associations between each of the posterior medians of the subject-level model parameters (β, *φ*, ρ) and the two DA baseline measures (blink rate, working memory compound score). Models with and without quadratic terms were compared for each proxy measure via LOO cross-validation. In all cases, the fit of the model that included a quadratic term was poorer, and none of the quadratic terms were significant (all p>.05; see *SI* for details). We also found no evidence for a linear association between drug-related differences of subject-level model parameters (β, *φ*, ρ) and DA baseline measures with the help of linear regression models (p>.05 for all regression slopes; see *SI*).

### Distinct brain networks orchestrate exploration and exploitation

Analysis of the imaging data proceeded in two steps. First, we examined our data for overall effects of exploration/exploitation as well as for model-based parametric effects of prediction error (PE), expected value and uncertainty. In a second step, we examined the neural basis of the drug-induced change in exploration. All fMRI results are reported at a threshold of p<.05, FWE-corrected for whole brain volume, unless stated otherwise. Analyses of drug-induced changes were instead based on small volume FWE correction (p<.05) for seven regions that have previously been associated with exploration: the left/right FPC and left/right IPS (Daw et al., 2006), as well as the dACC and left/right AI (Blanchard & Gershman, 2018). Regions used for small volume correction were defined by 10mm radius spheres centered at peak voxels reported in these previous studies (see *SI*).

In a first general linear model (GLM), differences in brain activity between exploratory and exploitative choices were modeled (at trial onset) across all participants and drug conditions, using the binary classification previously described by Daw et al. (2006). In accordance with previous work, the pattern of brain activity differed markedly between both types of choices (Figure 5).

**Figure 5.**
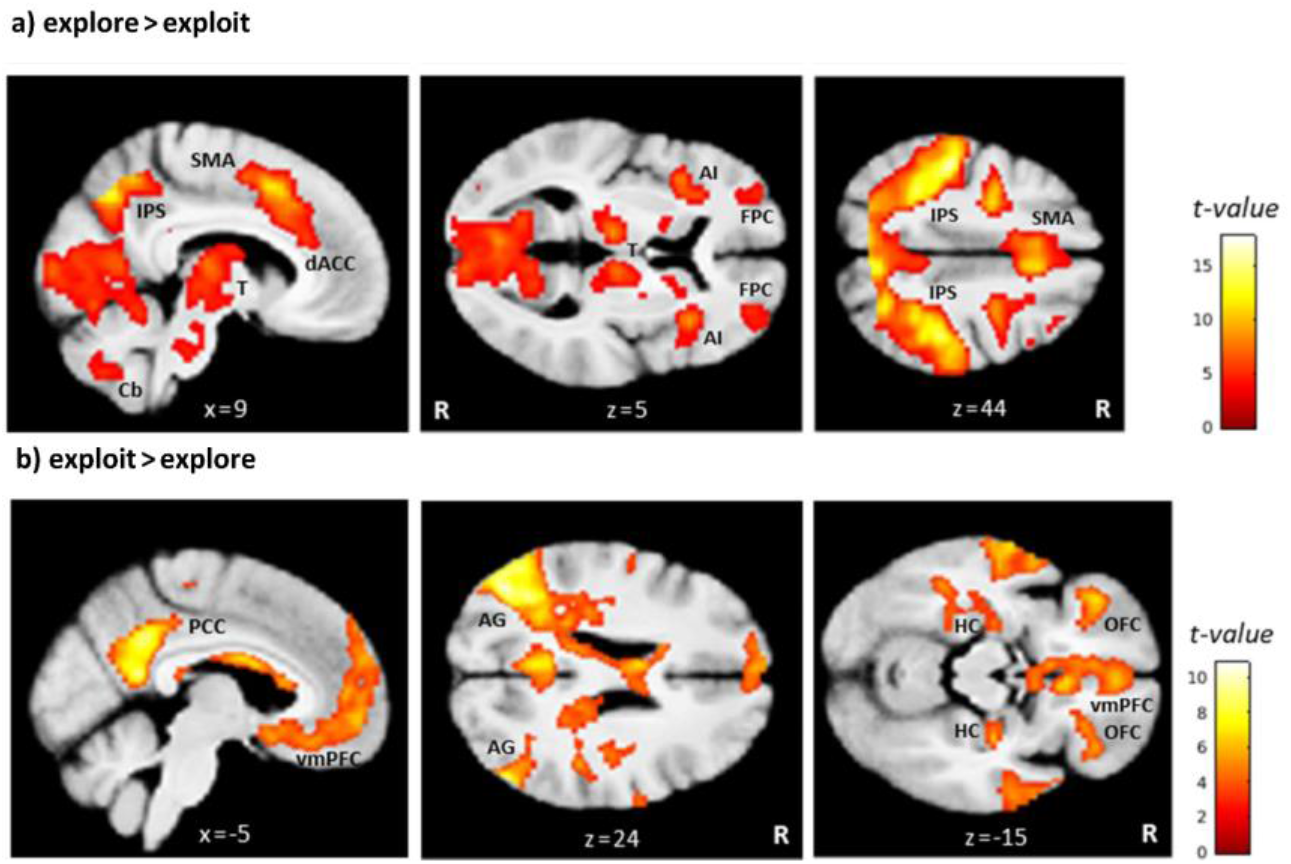
Brain regions differentially activated by exploratory and exploitative choices. Shown are statistical parametric maps (SPMs) for (a) the contrast explore > exploit and (b) the contrast exploit > explore over all drug conditions. AG: angular gyrus; AI: anterior insula; Cb: cerebellum; dACC: dorsal anterior cingulate cortex; FPC: frontopolar cortex; HC: hippocampus; IPS: intraparietal sulcus; vmPFC: ventromedial prefrontal cortex; OFC: orbitofrontal cortex; PCC: posterior cingulate cortex; SMA: supplementary motor area; T: thalamus. For visualization purposes thresholded at p < .001, uncorrected. R: right.

Replicating earlier findings (e.g. Daw et al., 2006; Addicott et al., 2014), exploration trials were associated with greater activation in bilateral frontopolar cortex (FPC; left: −42, 27, 27 mm; z=6.07; right: 39, 34, 28 mm; z=7.56), in a large cluster along the bilateral intraparietal sulcus (IPS; cluster peak at −48, −33, 52; z = 10.45), and in bilateral anterior insula (AI; left: −36, 15, 3 mm; z=6.69; right: 36, 20, 3 mm; z=6.87) as well as dorsal anterior cingulate cortex (dACC; cluster peak at 8, 12, 45 mm; z=8.47). Clusters within thalamus, cerebellum, and supplementary motor area also showed increased bilateral activation during exploration compared to exploitation.

In contrast, exploitative choices were associated with greater activation in the ventromedial prefrontal cortex (vmPFC; −2, 40, −10 mm; z=5.67) and in bilateral lateral orbitofrontal cortex (lOFC; left: −38, 34, −14 mm; z=5.81; right: 38, 36, −12 mm; z=5.02). Furthermore, greater activation during exploitative trials was also observed in a cluster spanning the left posterior cingulate cortex (PCC) and left precuneus (cluster peak at −6, −52, 15 mm; z=7.40), as well as in the angular gyrus (left: −42, −74, 34 mm; z=8.04; right: 52, −68, 28 mm; z=7.02), hippocampus (left: −24, −16, −15 mm; z=4.16; only at p < .001, uncorrected; right: 32, −16, −15 mm; z=5.09), and several clusters along the superior and middle temporal gyrus. A complete list of activations associated with explorative and exploitative choices can be found in *SI*.

The model-based PE was positively correlated with activity in bilateral ventral striatum (left: −16, 6, −12 mm; z=6.40; right: 16, 9, 10 mm; z=6.20), as shown in Figure 6.

**Figure 6.**
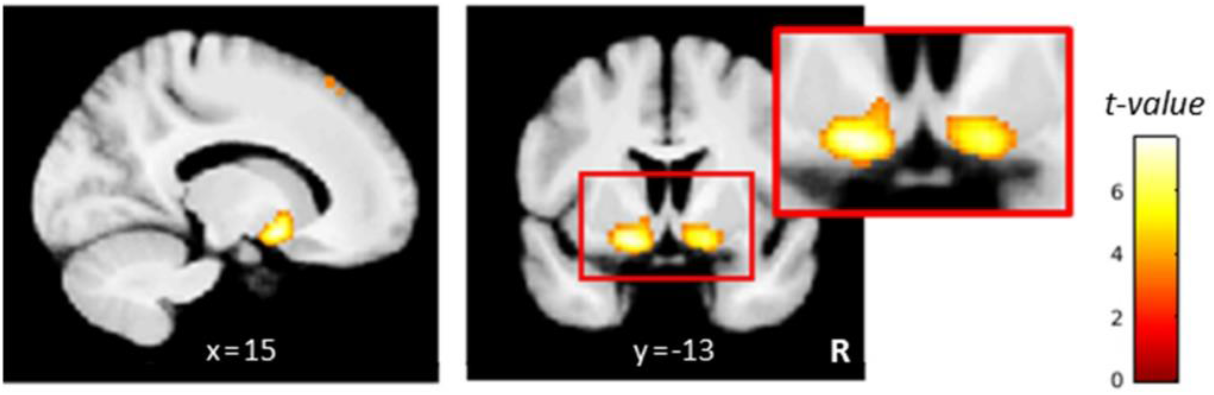
Striatal coding of the model-based prediction error (PE). Activity in the bilateral ventral striatum correlated positively with the PE signal. For visualization purposes thresholded at p < .001, uncorrected. R: right

In a second GLM we expanded upon the original trial classification of Daw et al. (2006) and included separate regressors for uncertainty-based and random exploration (both modeled at trial onset) to test for differential activation patterns of both exploration types. At p<0.05 FWE-corrected for whole-brain volume, the contrast uncertainty-based > random exploration yielded no supra threshold voxels across conditions or for any of the drug conditions alone. However, the reverse contrast (random >uncertainty-based exploration) yielded a small cluster of three voxels in the right FPC (32, 50, −8 mm; z=5.34) across conditions only. However, the number of trials was highly unequal for both exploration types with on average three times more random than uncertainty-based exploration trials per session. After exclusion of all sessions with ≤5 trials in the uncertainty-based exploration condition (8 out of 93 sessions), the frontopolar cluster was no longer significant at the whole-brain level. Furthermore, overlaying activation maps for uncertainty-based, random, and overall explorations (each contrasted against exploitation) showed a highly similar activation pattern for all three exploration conditions (see *SI*) that in each case included the same network (bilateral FPC, IPS, dACC, AI, and thalamus).

Finally, a third GLM was set up to examine parametric effects of exploitation and exploration (instead of the binary coding in the first GLM) based on two model-based quantities: the expected value 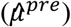 and uncertainty 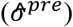 of the chosen bandit, both modeled at trial onset. While expected value of the chosen bandit was positively correlated with activity in a network of brain regions largely overlapping with the one for exploitative choices (see Figure 7a), uncertainty of the chosen bandit was positively correlated with activity in a network of brain regions largely overlapping with the one for exploratory choices (see Figure 7b).

**Figure 7.**
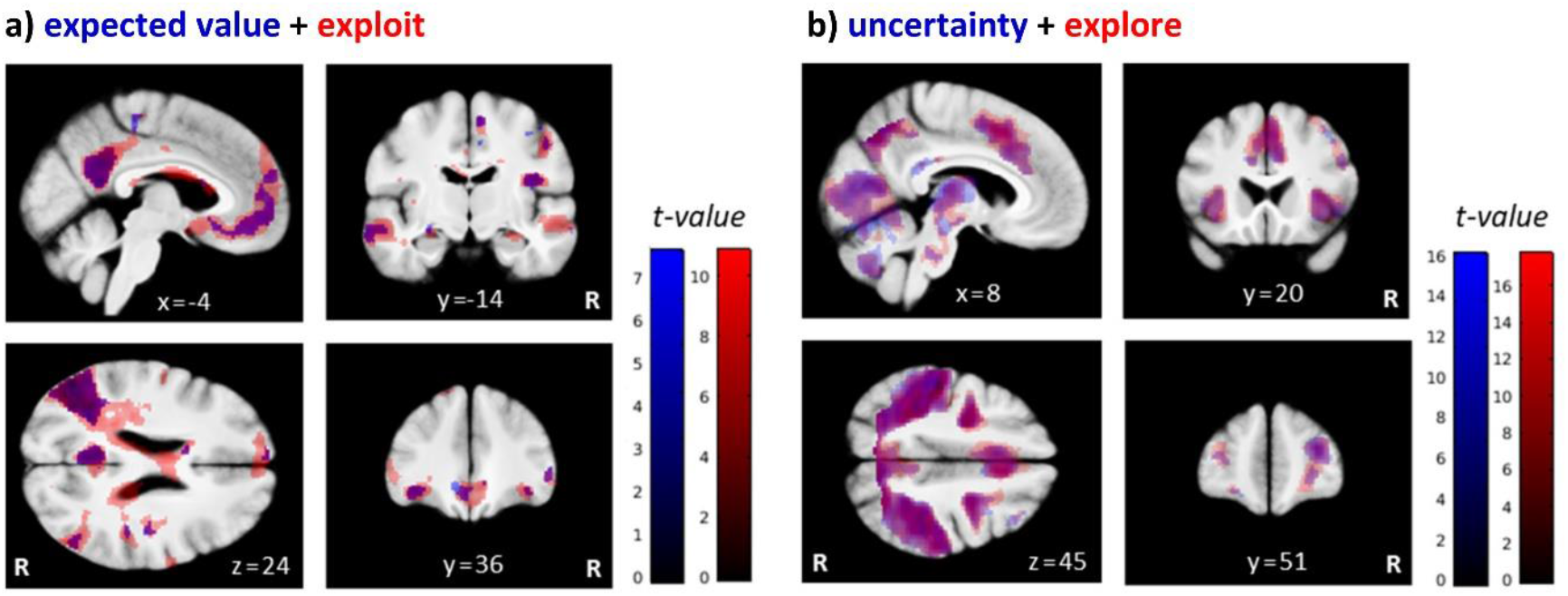
Neural coding of expected value 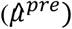 and uncertainty 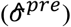. Overlay of the SPMs for (a) the parametric regressor expected value (in blue) and the contrast exploit > explore (“exploit” in red), and for (b) the parametric regressor uncertainty (in blue) and the contrast explore > exploit (“explore” in red), over all drug conditions. For visualization purposes: thresholded at p < .001, uncorrected. R: right.

### No evidence for a direct drug modulation of exploration/exploitation-related brain activation

To test for a main effect of drug on differential explore/exploit-related brain activation, we conducted repeated-measures ANOVAs on the second level contrasts explore vs. exploit of the first GLM. Surprisingly, we found no suprathreshold activations on the whole-brain level, nor in any of seven regions of interest (ROIs) with small volume correction applied (i.e. left/right FPC, left/right IPS, left/right AI, and dACC).

The same analyses were run for the contrasts from the second GLM, that is uncertainty-based exploration vs. exploit, random exploration vs. exploit, and random vs. uncertainty-based exploration. Again, we found no significant drug-related modulation of brain activation for these contrasts in any of our a priori ROIs.

In addition, we computed exploratory rmANOVAs for the four remaining regressors of the first GLM (trial onset, reward onset, PE, and outcome value), to explore other drug-related effects on brain activation. These analyses also revealed no suprathreshold activations on the whole-brain level.

DA drug effects on uncertainty-based exploration (*φ*) showed considerable variability between subjects both in terms of magnitude and direction (see Figure 4). Therefore, second-level regression analyses were performed for each drug pair, testing whether drug effects on exploration specific brain activity (first/second GLM contrasts for explore vs. exploit, uncertainty-based vs. exploit, and random vs. exploit) were linearly predicted by the drug effects on exploratory behavior (drug related differences of the subject-specific *φ* medians). However, none of these analyses revealed any suprathreshold effects on the whole brain level, nor in any of the seven a priori ROIs.

### L-dopa indirectly modulates exploration via reducing neural coding of overall uncertainty

Based on the null findings in the planned analyses, we reasoned that L-dopa might attenuate uncertainty-based exploration not by modulating brain activation for exploratory/exploitative choices in a direct manner, but rather by modulating neural computations that are involved in switching from exploitation to uncertainty-based exploration. Thus, L-dopa might delay the time point at which uncertainty-based exploration is triggered in response to accumulating uncertainty. Within our model, the overall (summed) uncertainty 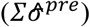 over all bandits is directly linked (via the exploration bonus parameter *φ*) to the probability to explore a previously unchosen bandit. Thus, 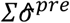 gradually increases during a series of exploitations but reduces abruptly when a bandit with high uncertainty is explored (see Figure 2f). We therefore included model-based overall uncertainty as a parametric regressor modeled at trial onset in a new GLM to reveal brain activation that tracks accumulating uncertainty during learning. The contrast images for this regressor were then used in a second-level random effects analysis with the factors drug condition and subject. In the placebo condition alone, no voxels survived whole-brain FWE correction (p < .05), but a more lenient threshold (p < .001, uncorrected) revealed activity in the bilateral dACC (cluster peak at −3, 21, 39 mm; z = 3.96), right anterior insula (42, 15, −6 mm; z = 3.46), and left posterior insula (PI) (−34, −20, 8 mm; z = 4.63) that was positively correlated with the overall uncertainty (Figure 8a). To test our exploratory hypothesis, we computed directed t-contrasts for L-dopa vs. placebo (placebo > L-dopa and L-dopa > placebo). While the contrast L-dopa > placebo yielded no suprathreshold activations, the reverse contrast (placebo > L-dopa) revealed a significant activation in the left PI (−34, −20, 8 mm; z = 5.05). At a reduced threshold (p < .001, uncorrected), left AI (−38, 6, 14 mm; z = 4.88) and bilateral dACC (left: −2, 36, 33 mm; z = 3.32; right: 4, 14, 28 mm; z = 3.41) showed a stronger correlation with the overall uncertainty under placebo compared to L-dopa (Figure 8b).

**Figure 8.**
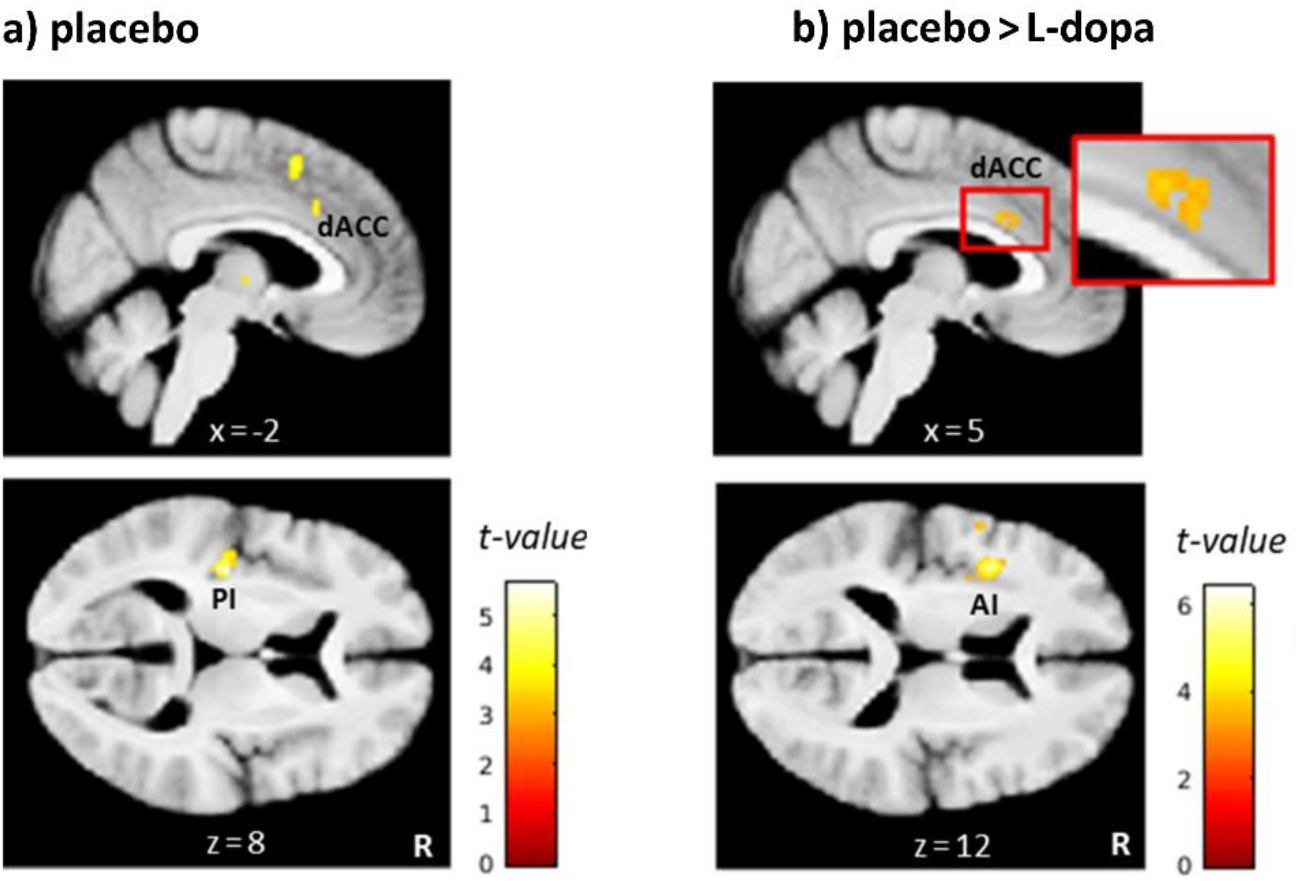
L-dopa effects on neural coding of overall uncertainty. (a) Regions in which activity correlated positively with the overall uncertainty in the placebo condition included the dorsal anterior cingulate cortex (dACC) and left posterior insula (PI). (b) Regions in which the correlation with overall uncertainty was reduced under L-dopa compared to placebo included the dACC and left anterior insula (AI). Thresholded at p < .001, uncorrected. R: right.

To directly link these exploratory findings with the behavioral L-dopa effect, we computed a second-level regression analysis on the overall uncertainty regressor for placebo > L-dopa. This regression analysis tested whether the L-dopa effects on uncertainty-related neural activation were linearly predicted by the L-dopa effects on uncertainty-based exploration, that is L-dopa-related differences of the subject-specific *φ* medians compared to placebo. However, this regression analysis yielded no suprathreshold activations.

## Discussion

We directly tested the causal role of DA in human explore/exploit behavior in a pharmacological, computational fMRI approach, using L-dopa (DA precursor) and haloperidol (DA antagonist) in a double-blind, placebo-controlled, counterbalanced, within-subjects design. Model comparison revealed that choice behavior was best accounted for by a novel extension of a Bayesian learning model (Daw et al. 2006), that included separate terms for uncertainty-based exploration and choice perseveration. Modeling revealed that uncertainty-based exploration was reduced under L-dopa compared to placebo and haloperidol. In contrast, no drug effects were observed on parameters capturing random exploration (β) or perseveration (ρ). Extensive additional analyses revealed no support for a modulation of DA drug effects by proxy measures of DA baseline (eye blink rate and working memory). On the neural level, exploration was associated with higher activity in the FPC, IPS, dACC, and insula, whereas exploitation showed higher activity in the vmPFC, OFC, PCC, precuneus, angular gyrus, and hippocampus, replicating previous studies (Daw et al., 2006; Addicott et al., 2014; Blanchard & Gershman, 2018). Surprisingly, no drug effects were found for these effects, nor on striatal reward prediction error signaling. However, an exploratory model-based analysis revealed that L-dopa reduced insular and dACC activity associated with overall (summed) uncertainty.

### Uncertainty-based exploration and perseveration

We examined two learning rules (Delta rule vs. Bayesian learner) and three choice rules resulting in a total of six computational models. Model comparison revealed that the Bayesian learning model (Kalman Filter) outperformed the Delta rule for each of the choice rules. Although both learning rules are based on the same error-driven learning principle, the Bayesian learner assumes that subjects additionally track the variance (uncertainty) of reward expectation and adjust the learning rate from trial to trial according to the current level of uncertainty - learning is high when reward predictions are uncertain (i.e. during exploration), but decreases when predictions become more accurate (i.e. during exploitation).

The three choice rules implemented within the different models were (1) a standard softmax (only capturing ‘random exploration’), (2) softmax with an additional parameter modeling uncertainty-based uncertainty-based exploration via an exploration bonus parameter, and (3) softmax with parameters for both uncertainty-based exploration and perseveration. For both learning rules the full model including uncertainty-based exploration and perseveration accounted for the data best. Inclusion of an exploration bonus parameter (φ) improved model fit, a finding consistent with previous research that showed that humans use both random and uncertainty-based exploration strategies (Frank et al., 2009; Cavanagh et al., 2012; Krueger et al., 2017; Payzan LeNestour & Bossaerts, 2012; Wilson et al., 2014; Gershman, 2018;). Random and uncertainty-based exploration were implemented via separate parameters, the first by adding stochasticity to the action selection process via softmax (Thrun, 1992; Daw et al., 2006; Beeler et al., 2010; Gershman, 2018) and the latter via an exploration bonus parameter (e.g. Dayan & Sejnowski, 1996; Daw et al., 2006). In this scheme, “uncertainty” or “information” biases choices towards more uncertain/informative options by increasing their value.

However, it has been argued before that perseveration, that is, repeating choices regardless of value (Schönberg et al., 2007; Brough et al., 2008; Rutledge et al., 2009; Worthy et al., 2013) can negatively impact estimates of uncertainty-based exploration (Daw et al., 2006, Bardre et al., 2012; Payzan-LeNestour & Bossaerts, 2012; Wilson et al., 2014), leading to smaller or even negative exploration bonus effects. To address this issue, we included a perseveration term in the exploration bonus model (Daw et al., 2006). This not only improved model fit, but also substantially increased estimates of uncertainty-based exploration and reduced the number of subjects showing a negative *φ* estimate.

### Behavioral DA drug effects

Overall, our finding that pharmacological manipulation of the DA system impacts the exploration / exploitation trade-off is in line with previous animal and human studies (e.g. Frank et al., 2009; Beeler et al., 2010; Blanco et al., 2015). However, the observed pattern of drug effects did not match our initial hypothesis, according to which both random and uncertainty-based exploration were expected to *increase* under L-dopa vs. placebo and *decrease* under haloperidol vs. placebo.

#### L-dopa

L-dopa administration reduced uncertainty-based exploration (*φ*) compared to placebo, while random exploration (β) was unaffected. In the model, this could reflect (1) a reduced tendency for uncertainty-based exploration and/or (2) an increased tendency for value-driven exploitation. Accordingly, when classifying all choices per subject into exploitations, uncertainty-based exploration and random exploration, L-dopa was found to reduce the percentage of uncertainty-based but not random exploration compared to placebo across subjects, and marginally increase the percentage of exploitations (see SI).

Since on the neural level, L-dopa is well known to increase DA transmission compared to placebo, these findings suggest that increased DA might increase exploitation vs. uncertainty-based exploration. This interpretation is in line with several studies showing that striatal DA drives reinforcement learning and exploitation (e.g. Schultz et al., 1999; Pessiglione et al., 2006; Frank et al., 2004, 2009). Furthermore, the observed L-dopa effect also informs an ongoing debate regarding DA contributions to model-based (goal-directed) vs. model-free (habitual) components of reinforcement learning and decision-making. In the Bayesian learning model of the bandit task, uncertainty-based exploration relies on an internal model of the environment (the Gaussian random walks). In the two-step task (Daw et al., 2011), a prominent task to dissociate model-based from model-free contributions to behavior, an earlier study reported L-dopa to increase model-based learning (Wunderlich et al., 2012). In contrast, a recent study in a considerably larger sample (N=64) reported L-dopa to decrease model-free learning, while increases in model-based learning where restricted to a subset of participants (Krömer et al., 2019). Furthermore, Deserno et al. (2015) reported a positive association between striatal presynaptic DA levels and the degree of model-based control in the two-step task. While these findings support the general idea that striatal DA might contribute to model-based decision-making, it contrasts with our finding of an L-dopa induced *reduction* of (presumably model-based) uncertainty-based exploration. However, uncertainty-based exploration and model-based control in sequential decision tasks might also rely on dissociable mechanisms. For example, while the hippocampus is implicated in model-based planning in the two-step task (Vikbladh, Meager, King et al. 2019), in our data hippocampal activation was more pronounced during exploitation. This discrepancy might also relate to differences in cognitive load that is arguably higher in the two-step task due its sequential structure. In the more complex two-step task, participants might benefit from an increase in DA transmission enabling them to rely more on cognitively demanding model-based strategies and less on computationally less expensive model-free mechanisms.

In contrast, studies that focused on prefrontal DA mostly found that DA promotes exploration (Frank et al., 2009; Blanco et al., 2015; Kayser et al., 2015). This inconsistency is likely resolved when considering the locus of action of L-Dopa, which likely increases DA transmission most prominently within striatum, and to a much lesser degree in PFC (Carey, Dai et al., 1995; Cools et al., 2006). In rodents, L-dopa administration generates about 50 times more DA in striatum than cortex (Carey, Dai et al., 1995). It is further assumed that L-dopa primarily affects phasic rather than tonic striatal DA activity in healthy individuals. This is supported by PET studies in humans (Floel et al., 2008; Black et al., 2015) that assessed striatal DA release with the radioligand [^11^C]raclopride (RAC), which binds to D2/D3-like receptors but competes with endogenous DA. While L-dopa (vs. placebo) administration produced no measurable increase in baseline (tonic) striatal DA release under resting conditions (Black et al., 2015; Floel et al., 2008), it induced a significant increase in task-evoked (phasic) striatal DA release during motor training (Floel et al., 2008). This increased phasic DA release was associated with improved learning under L-dopa compared to placebo, presumably by enhancing the reinforcing effect of positive feedback during learning (Frank et al., 2004; Cox et al. 2015; Mathar et al., 2017).

Mechanistically, exogenous L-dopa is assumed to be taken up by nigro-striatal dopaminergic nerve terminals, converted into DA, stored in synaptic vesicles and co-released with endogenous DA upon neural excitation (Horne et al., 1984; Breitenstein et al., 2006; Floel et al., 2008). Thus, L-dopa may “stock up” presynaptic DA stores in healthy subjects, in which DA catabolism and storage capacity are sufficient to prevent exogenous DA from being released under resting conditions, i.e. during tonic firing. In individuals suffering from depleted DA levels, such as Parkinson’s disease patients, L-dopa may “replenish” DA stores and boost both tonic and phasic DA transmission. Our finding that L-dopa decreased uncertainty-based exploration is in line with the assumption that L-dopa primarily boosts phasic but not tonic DA transmission in healthy individuals. Studies that specifically examined effects of tonic DA modulation on exploration/exploitation suggest that elevated tonic DA levels in striatum increase rather than decrease exploration (Beeler et al., 2010; Costa et al., 2014). Costa et al. (2014) found that monkeys with elevated striatal tonic DA levels due to DAT blockade showed increased uncertainty-based exploration. One explanation for this opposite effect is that increased tonic DA levels attenuate phasic transmission via inhibitory presynaptic D2 autoreceptors (Grace, 1991; Cools, 2006; Ford, 2014). DAT knockdown mice were shown to exhibit not only elevated tonic striatal DA levels, but also a clear (ca. 75 %) reduction in the amplitude of phasic striatal DA release (Zhuang et al., 2001). In line with studies on temporal discounting (Pine et al., 2010), L-dopa might strengthen positive reinforcing effects of immediate rewards via increased phasic striatal DA release, fostering both impulsive and exploitative choice behavior (Kobayashi & Schultz, 2008; Schultz, 2010; Pine et al., 2010).

Importantly, the striatum is densely interconnected with frontal cortices including both bottom-up (striatum➔PFC) and top-down (PFC➔striatum) projections (Badre & Nee, 2018; Haber & Knutson, 2010). Prefrontal DA has been repeatedly related to uncertainty-based exploration (Frank et al., 2009; Blanco et al., 2015; Kayser et al., 2015). Human brain stimulation experiments suggest a causal role of the fronto-polar PFC in uncertainty-based exploration (Raja Beharelle et al., 2015; Zajkowski et al., 2017). In light of these previous results, our finding of reduced uncertainty-based exploration under L-dopa might also relate to a shift in the balance between striatal and prefrontal control over the balance between exploration and exploitation. Uncertainty-based exploration might be implemented via a fronto-striatal top-down control mechanism (Raja Beharelle et al., 2015; Zajkowski et al., 2017). In such a model, FPC tracks the relative uncertainty of choice options and modulates striatal DA transmission via top-down connections to override exploitation and facilitate exploration. This is supported by studies reporting an inverse relationship between frontal and striatal DA activity (Akil et al., 2003; Meyer-Lindenberg et al., 2005; Cools & D’Esposito, 2011). Striatal DA transmission might also modulate frontal DA activity via bottom-up interference (Kellendonk et al., 2006; Simpson, Kellendonk, & Kandel, 2010; Duvarci et al., 2018). In support of this idea, in humans, striatal D2/D3 receptor availability was positively correlated with reward-related striatal activity, but negatively with risk-related prefrontal activity and risky choice behavior (Kohno et al., 2015). In such a model, an L-dopa-induced increase in phasic DA transmission may also attenuate exploration via an interference with prefrontal (dopaminergic) processing.

#### Haloperidol

Contrary to our hypothesis, we did not observe a significant modulation of exploration/exploitation under haloperidol. Computational modeling revealed no changes in the group-level mean parameters for uncertainty-based exploration (φ), random exploration (β) or perseveration. However, upon closer inspection, haloperidol may show a complex effect on the uncertainty-based exploration parameter *φ*, with a tendency to increase *φ* in low-φ subjects (i.e. low *φ* at placebo) and to decrease *φ* in high-φ subjects. This is also supported by the finding that haloperidol showed a tendency to decrease the group-level variance of *φ* (Figure 3c).

Haloperidol is a potent D2 receptor antagonist. Thus, the absence of a clear effect on exploration and/or exploitation is at first glance somewhat puzzling. We predicted haloperidol to attenuate tonic DA signaling and reduce directed and random exploration. However, alternatively one could have predicted haloperidol to reduce phasic signaling within striatum and attenuate reward exploitation and thus increase exploration (Pessiglione et al., 2006; Pleger et al., 2009; Eisenegger et al., 2014). For example, Pessiglione et al. (2006) reported that, compared to L-dopa, 1mg of haloperidol reduced reward exploitation and reward PE signaling within striatum in a reinforcement learning task. We did not find evidence for this alternative hypothesis, neither in the behavioral nor in the neural data. In contrast to Pessiglione et al. (2006), no drug effects on striatal PE coding were observed, neither for L-Dopa nor Haloperidol.

Importantly, numerous studies also found opposite effects of single doses of haloperidol, and the directionality of effects might depend on the dosage. Low doses of D2 antagonists can stimulate DA signaling which contrasts with the antidopaminergic effects observed under chronic and high-dose treatment (Starke et al., 1989; Frank & O’Reilly, 2006; Knutson & Gibbs, 2007). In addition to postsynaptic receptors, D2 agents can also act on presynaptic auto-receptors. While blockade of postsynaptic receptors reduces DA signaling, blockade of presynaptic auto-receptors may stimulate DA transmission due to a reduced feedback inhibition of DA release (Schmitz et al., 2003; Frank & O’Reilly, 2006; Ford, 2014). Lower doses of D2 antagonists might primarily act on presynaptic receptors and thus increase phasic DA release (Youngren, 1999; Westerink et al., 2001; Schwerdt et al., 2017) and related striatal blood volume (Schwarz et al., 2004; Chen et al., 2005; Handley et al., 2013). This could also explain paradoxical drug effects of D2 antagonists observed in several human studies. With the same dose of 2mg as in our study, Frank & O’Reilly (2006) found haloperidol to enhance learning from positive feedback whereas it diminished learning from negative feedback. Similar effects were observed with a low dose of amisulpride, a different D2 antagonist (Jocham, Klein, & Ullsperger, 2011). Other studies that used low doses of haloperidol found no consistent drug effect or only effects in subgroups of participants (Frank & O’Reilly, 2006; Pine et al. 2010). This inconsistency might be explained by a dependency of drug-effects on the individual baseline level of D2 receptor availability (Frank & O’Reilly, 2006; Kirsch et al., 2006; Eisenegger et al., 2014). To address this issue, several studies used measures of working memory capacity as an index of striatal DA transmission (Frank & O’Reilly, 2006; Gibbs & D’Esposito, 2005; van der Schaaf et al., 2014), a discussion that we return to below.

Taken together, our null finding regarding the effect of haloperidol might be related to several factors, including a dosage that was too low to act on postsynaptic receptors (but see Pessiglione et al., 2006), and a potential baseline dependency that gave rise to a complex non-linear pattern of drug effects. However, we also examined two proxy measures for striatal DA transmission (eye blink rate and working memory capacity), both of which showed no modulation of drug-effects (see below).

### fMRI findings

#### Neural correlates of exploration and exploitation

Consistent with previous research, the pattern of brain activity differed markedly between exploration and exploitation: Exploratory choices were associated with higher activity in the FPC, IPS, dACC, and AI, replicating previous human fMRI studies (Daw et al., 2006; Addicott et al., 2014; Laureiro-Martínez et al., 2014, 2015). It has been suggested that the FPC may track information relevant for exploratory decisions, such as the expected reward and uncertainty of unchosen choice options, and trigger a behavioral switch from an exploitative to an exploratory mode whenever the accumulated evidence supports such a switch (Boorman et al., 2009, 2011; Badre et al., 2012; Cavanagh et al., 2012). However, this interpretation would predict that FPC activation should be greater for directed vs. random exploration, an effect that was not observed in the present study when accounting for the different trial numbers for the two types of choices (see discussion below). The IPS, in contrast, has been suggested to serve as an interface between frontal areas and motor output areas, initiating behavioral responses to implement exploratory actions (Daw et al., 2006; Boorman et al., 2009; Laureiro-Martínez et al., 2015). The dACC and AI, on the other hand, are thought to form a salience network involved in detecting and orienting towards salient stimuli (Menon, 2015; Uddin, 2015), which may also subserve attentional and behavioral switching from exploitation to exploration. Aside from these regions, bilateral thalamus, cerebellum, and supplementary motor area showed greater activation during exploration compared to exploitation, findings mostly consistent with previous fMRI studies (Daw et al., 2006; Addicott et al., 2014; Laureiro-Martínez et al., 2014, 2015).

Exploitative choices were associated with greater activation in vmPFC and OFC, again replicating previous work (Laureiro-Martínez et al., 2014, 2015). Both regions are implicated in coding a subjective value of attainable goods (Levy & Glimcher, 2012). Thus, vmPFC and OFC might foster exploitation based on computing subjective values of decision options (Kringelbach & Rolls, 2004; O’Doherty, 2004, 2011; Peters & Büchel, 2010; Bartra et al., 2013).

In addition, greater activation during exploitation was also observed in the PCC, angular gyrus, precuneus, and hippocampus, partly replicating the results of earlier studies (Addicott et al., 2014; Laureiro-Martínez et al., 2014, 2015). Together with the medial PFC, these regions are hypothesized to form a large-scale brain system referred to as the “default mode network” (DMN; Raichle et al., 2001; Andrews-Hanna, Smallwood, & Spreng, 2014). Thus, activity within these regions during exploitation may also relate to a reduced cognitive and attentional demand during exploitation compared to exploration. Interestingly, the salience network (dACC and AI) has been proposed to play a key role in switching brain activity from introspective functions of the DMN to externally focused, task-based functions (Bressler & Menon, 2010; Menon, 2015), consistent with our finding that regions of this salience network showed higher activity during exploration (see above). Alternatively, activation of these regions may also subserve more specific exploitation-related functions. The angular gyrus is also heavily implicated in number monitoring (Göbel et al., 2001) and thus may monitor reward values during exploitation (see Addicott et al., 2014). The PCC is considered to be part of the brain’s valuation system and hence might also be involved in encoding reward-related information during exploitation (Lebreton et al., 2009; Bartra et al., 2013; Grueschow et al., 2015). Moreover, increased hippocampal activity during exploitation may reflect processes of episodic memory retrieval involved in reward-based decision making (Bornstein et al., 2017). A further characterization of the hypothesized functions of subregions of the exploitation- and exploration networks naturally requires direct experimental tests in the future.

In additional analyses, we examined parametric effects of several model-based quantities. The reward PE signal was found to positively correlate with activity in the bilateral ventral striatum, consistent with numerous previous studies (O’Doherty et al., 2003, 2004; Hare et al., 2008; Gläscher, Daw & O’Doherty., 2010). We also examined two parametric predictors directly linked to explore/exploit behavior: the expected value 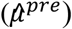 and uncertainty 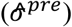 of the chosen bandit. Parametric neural effects for expected reward and uncertainty largely overlapped with networks for exploitation and exploration from the categorical analysis (see *SI* and Figure 7). This confirms that the neural effects are robust to different parameterizations of the respective processes.

In a novel analysis approach that extended previous work (Daw et al., 2006), we examined whether different neural substrates for random and uncertainty-based exploration could be identified. Across subjects and drug conditions, random exploration was associated with greater activity in a small region of the right FPC compared to uncertainty-based exploration. Yet, after accounting for the unequal number of trials in both exploration conditions, the fronto-polar activation differences between uncertainty-based and random exploration where no longer significant. This may partly be related to the definition of uncertainty-based exploration utilized here (choosing the bandit with highest exploration bonus). Since the exploration bonus influences choice behavior parametrically, also choices classified as random exploration might contain signatures of uncertainty-based exploration. Finally, while the bandit task might resemble real-world exploration behavior and how it evolves over longer time frames quite well, other tasks might allow for a clearer separation of random and uncertainty-based exploration (Wilson et al., 2014).

In contrast to earlier models (Aston-Jones & Cohen, 2005; Humphries et al., 2012; Mandali et al., 2015), recent accounts propose that random exploration might also be driven by uncertainty (Gershman, 2018). Both exploration types might thus depend on similar structures tracking uncertainty. Mansouri et al. (2017) recently proposed a functional model of the human FPC in which distinct subregions of the FPC play different functional roles in exploration. In this model, lateral FPC is involved in uncertainty-based exploration, which entails an online tracking of relevant choice alternatives in order to potentially re-engage one of these alternatives as replacement for the currently exploited strategy. In contrast, medial FPC is involved in random exploration, for which only the ongoing strategy is monitored to potentially redistribute cognitive resources away from this strategy when it is deemed irrelevant. Hence, our finding of a small FPC subregion with somewhat higher activity during random than uncertainty-based exploration would be consistent with such a functional model.

Notably, the cognitive model applied here nests the exploration bonus within the softmax function. Consequently, random exploration in our modeling framework always occurs relative to the sum of expected value, exploration bonus and perseveration bonus. This formalization has the advantage that bonuses can be directly interpreted in terms of reward value units (points) allowing a quantitative comparison of these factors. However, the idea that random exploration might also be driven by uncertainty (see above; Gershman, 2018) is not captured by this model.

#### Neural DA drug effects

We found no drug effects on exploration- or exploitation-related brain activity, nor on the neural correlates of reward PE signals. For the L-dopa condition, the null results for the planned explore/exploit comparisons are particularly surprising, given the clear behavioral L-dopa effect on uncertainty-based exploration. We hypothesized DA drug effects on exploratory behavior to be associated with changes in the activity of brain regions implicated in exploratory choices, in particular the FPC and IPS. This was not supported by the fMRI data. Alternatively (or additionally), the observed L-dopa effect on explore/exploit behavior could also be due to an enhanced phasic DA release and PE signaling in the striatum (see above). In such a model, L-dopa would be expected to increase the magnitude of the striatal reward PE signal, as previously shown by Pessiglione et al. (2006). However, this effect could not be replicated in the current study. This could be due to a range of factors. Pessiglione et al. (2006) used a between-subjects design with a simpler task, while we adopted a within-subjects design with a more complex and volatile learning environment. Sample sizes also varied considerably (n=13 per group in Pessiglione et al. (2006), n=31 in the present study). A recent study (Kroemer et al. (2019)) also did not observed a modulatory effect of L-dopa on PE coding in a sequential reinforcement learning task in an even larger sample (N=65).

Several additional factors might have contributed to the failure to detect L-dopa effects on the neural correlates of explore/exploit decisions or the reward PE. First, this failure may simply be due to a lack of statistical power provided by the modest sample size of 31 subjects (Button et al., 2013; Turner, et al., 2018). However, as noted above, previous studies used a much smaller sample size, and in our design power was increased due to the within-subjects design. Furthermore, the timing of the drug administration might also be a crucial factor to consider. L-dopa has a plasma half-life of 60 to 90 min and reaches peak plasma concentrations (tmax) about 30 to 60 min after oral ingestion (Baruzzi et al., 1987; Keller et al., 2011; Iwaki et al., 2015). The time schedule of the current experiment was adjusted to this tmax: The bandit task started 30 min and ended 80 min after L-dopa administration, such that peak plasma concentrations were reached approximately halfway through the task. However, it is also conceivable that L-dopa effects on phasic DA activity might peak with some additional delay, considering that L-dopa needs to pass the blood-brain barrier (by active transport), be converted to DA and packaged into synaptic vesicles to contribute to phasic DA signaling. Also, tmax is usually measured after over-night fasting, whereas subjects in the current study were not fasted to avoid fasting-related effects on behavior. This may have delayed L-dopa effects in this study due to slower gastric emptying and increased competition between L-dopa and dietary amino acids for active transport across the intestine and blood-brain barrier (Baruzzi et al., 1987; Contin & Martinelli, 2010; Wang et al., 2017). Notably, the study of Pessiglione et al. (2006), in which L-dopa was found to increase striatal PE signaling, followed a slightly different time schedule for drug administration. In that study, the reinforcement learning task started one hour after L-dopa administration, which could have been more suitable to capture the time interval in which L-dopa exerts its maximal neural and behavioral effects. However, such considerations fall short in explaining the clear behavioral effect of L-Dopa that was observed in the present study. It should also be noted that behavioral drug-effects in Pessiglione et al. (2006) where only observed in the direct comparison of the haloperidol and L-Dopa groups, and not in comparison to placebo.

Finally, the BOLD signal obviously does not directly reflect DA release, and the precise physiological relationship between DA release and BOLD signal is still to be revealed (Knutson & Gibbs, 2007; Brocka et al., 2018). Based on pharmacological MRI studies, it has been suggested that striatal DA release may increase the BOLD signal via a D1-dependent mechanism, according to which D1 receptor activation changes the postsynaptic membrane potential and engages metabolic processes, which in turn lead to increased oxygen utilization followed by an elevated local BOLD response (Knutson & Gibbs, 2007). However, a recent optogenetic study in rats suggests that canonical BOLD responses in the reward system may mainly represent the activity of non-dopaminergic neurons, such as glutamatergic projecting neurons (Brocka et al., 2018). Thus, it is also conceivable that L-dopa might have enhanced striatal DA release to some degree without triggering a (detectable) BOLD signal change.

For the haloperidol condition, the null findings on the neural level are less surprising, given the lack of a consistent behavioral effect across subjects. As discussed above, it can be assumed that the low dose (2 mg) of haloperidol used in this study exerted a mixture of presynaptic (DA-stimulating) and postsynaptic (DA-antagonizing) effects across subjects, potentially explaining why no overall haloperidol effects were found on the behavioral and neural level. Similarly, Pine et al. (2010) also did not observe significant effects of haloperidol (1.5mg) on reward-related striatal activity or choice behavior. On the other hand, Pessiglione et al. (2006) reported that haloperidol (1 mg) reduced the magnitude of the striatal reward PE signal and exploitative behavior relative to L-dopa. Yet, as mentioned above, haloperidol effects were reported only relative to the L-dopa condition, as the placebo condition was not double-blinded. Visual inspection of the behavioral data of Pessiglione et al. (2006) suggests that L-Dopa (rather than haloperidol) drove the observed effects.

Future studies should consider using higher doses of haloperidol to achieve more consistent antidopaminergic effects from postsynaptic D2 receptor blockade across subjects, or other DA antagonists with a lower side effect profile.

### L-dopa attenuates neural tracking of overall uncertainty

We reasoned that L-dopa may have reduced exploration not by affecting the neural signatures of categorical explore/exploit decisions, or the reward PE, but instead by modulating some other aspect of DA-dependent brain function involved in the explore/exploit trade-off. Specifically, we reasoned post-hoc that L-dopa may have affected the neural correlates involved in behavioral switching from exploitation to exploration in response to accumulating uncertainty. Thus, L-dopa might delay the time point at which uncertainty-based exploration is triggered, resulting in less uncertainty-based exploration trials over time. We examined this alternative hypothesis with an additional model-based fMRI analysis, in which a trial-by-trial estimate for overall uncertainty (summed standard deviation over all bandits), was used as a parametric regressor in the GLM. Activity in the bilateral insula and dACC positively correlated with the overall uncertainty in the placebo condition, suggesting that these regions may either track the overall uncertainty directly or encode an affective or motivational state that increases with accumulating uncertainty. Insula and dACC thus may trigger exploration under conditions of high overall uncertainty, e.g. by facilitating switching between the currently exploited option and salient, more uncertain choice alternatives (Kolling et al., 2012, p. 97; Laureiro-Martínez et al., 2015).

Indeed, numerous studies have found greater activation in these regions during decision-making under uncertainty, and have implicated both regions in encoding outcome uncertainty or risk (Huettel et al., 2005; Preuschoff et al., 2006, 2008; Christopoulos et al., 2009; Singer et al., 2009; Bach & Dolan, 2012; Dreher, 2013). The insula is also considered to play a key role for integrating interoceptive signals about bodily states into conscious feelings, such as urgency, that can influence decision-making under risk and uncertainty (Craig, 2002, 2009; Critchley, 2005; Naqvi & Bechara, 2009; Singer et al., 2009; Xue et al., 2010). The ACC is known to play a crucial role in monitoring response conflict, which should increase with the overall uncertainty, and in triggering attentional and behavioral changes serving to reduce future conflict (Botvinick et al., 2004; Kerns et al., 2004; van Veen & Carter, 2002). Finally, both the ACC and insula project to the striatum (Kunishio & Haber, 1994; Haber & Knutson, 2010) where they modulate striatal reward signals and reward-related behavior (Walton et al., 2006; Behrens et al., 2007; Botvinick, et al., 2009; Shenhav et al., 2013). Accordingly, uncertainty-related activity in the insula and ACC, as observed in the current study, could directly interfere with the subcortical DA substrates of value-driven choice behavior to facilitate switching between exploitation and exploration. Taken together, these findings support the idea that both the insula and dACC are tightly involved in triggering exploration under circumstances of high overall uncertainty. The insula might signal an urge to explore that increases with accumulating uncertainty, whereas dACC might subserve cognitive control processes that guide attention and behavioral responses towards salient, uncertain choice alternatives.

Importantly, we found that L-dopa reduced uncertainty-related activity in the insula and dACC compared to placebo, potentially explaining why participants showed less uncertainty-based exploration under L-dopa. Expression of D1 and D2 receptors is much higher in striatum than in insula and ACC (Hall et al., 1994; Hurd et al., 2001), and L-dopa primarily exerts its effects within striatum (see discussion above). Hence, it seems more likely that L-dopa affected uncertainty-related activity in the insula and ACC indirectly by modulating DA transmission on the striatal level. More specifically, L-dopa might have modulated striatal processing of reward uncertainty (Preuschoff et al., 2006; Schultz et al., 2008) that subsequently is transmitted to cortical structures for integration with other decision parameters to guide explore/exploit behavior (Kennerley et al., 2006; Rushworth & Behrens, 2008; Haber & Knutson, 2010; Shenhav et al., 2013). Taken together, these findings suggest an interesting possibility regarding how L-dopa might have affected explore/exploit behavior. To further test these ideas, future studies should more closely examine the role of the insula and ACC in triggering exploration in response to accumulating uncertainty and further investigate how frontal and/or striatal DA transmission might modulate this process.

### Modulation of DA drug effects by DA baseline

One additional aim of this study was to test whether DA drug effects on explore/exploit behavior are modulated by individual DA baseline activity (as indexed by spontaneous blink rate and working memory capacity) as predicted by the inverted-u-shape hypothesis of DA (Cools & D’Esposito, 2011). However, we found no evidence for such a relationship between the DA baseline measures and DA drug effects on explore/exploit behavior in our data. This might be due to several reasons. Despite its popularity, the inverted-U-hypothesis is remains relatively vague regarding the specific DA functions and cognitive domains it applies to, and is therefore difficult to test and falsify. Animal studies suggest that it specifically describes the relationship between prefrontal D1 receptor activity and working memory performance, whereas the relation between other aspects of DA action and cognitive functions may follow different functions (Floresco & Magyar, 2006; Floresco, 2013).

It remains unclear how to construe the exact shape and turning point (optimum) of the inverted-U-shape function, since these aspects may vary across tasks, cognitive functions, and individuals (see Cools et al., 2009; Cools & D’Esposito, 2011; Wiegand et al., 2016). Obviously, our study also has limitations that exacerbate comprehensive testing of this more complex hypothesis. First, the sample size of 31 participants may simply be too small to detect an inverted quadratic relationship between proxies for DA baseline and drug effects on explore/exploit behavior. It has been argued (Slagter et al., 2012) that healthy subjects may display only a relatively restricted range in baseline DA levels during resting conditions, making it more difficult to observe inverted-u-shape effects in these samples. Blink rate values were strongly left-skewed across subjects, with only relatively few subjects with high blink rates, supporting this idea. The failure to observe an inverted-U-shaped effect in the current study might also be due to poor DA proxy measures. While the spontaneous blink rate has been extensively investigated as a proxy for DA function in animals and humans, many studies have also yielded conflicting or inconclusive results (Jongkees & Colzato, 2016; Dang et al., 2017; Sescousse et al., 2018). For working memory capacity, the available evidence is even more limited than for the blink rate, as fewer studies have used this measure as a DA proxy. While some studies report differential DA drug effects in relation to individual working memory capacity, the directionality of these effects differs between studies (Kimberg et al, 1997; Kimberg & D’Esposito, 2003; Gibbs & D’Esposito, 2005, 2006). Moreover, for both proxy measures, it is not clear which specific aspect of DA function they might index, and what the underlying mechanism is. Both measures might reflect aspects of striatal DA function, such as striatal D2 receptor availability (Groman et al., 2014; Jongkees & Colzato, 2016) and/or striatal DA synthesis capacity (Cools et al., 2008; Landau et al., 2009). In contrast, the inverted-U-shaped hypothesis predominantly relates to D1 receptor function in PFC. In conclusion, the DA proxies used in this study may have failed to validly measure baseline DA function, or specifically measured an aspect of DA function which was not predictive for the behavioral outcome measure under study.

### Limitations

In addition to the limitations discussed above, the moderate sample size of 31 subjects in this within-subjects manipulation study may have contributed to the absence of haloperidol effects and the absence of drug-associated differences in categorical contrasts of explore/exploit trials in the fMRI analysis. Power is an even greater issue for the hypothesized inverted-U-shape hypothesis (see above). A second limitation relates to the fact that while the applied pharmacological fMRI approach can examine DA drug effects on the BOLD signal, it remains unclear which effects directly reflect local changes in DA signaling, and which reflect downstream effects that may also involve other neurotransmitter systems (Schrantee & Reneman, 2014). Needless to say, the BOLD signal provides an indirect index of blood oxygenation rather than a direct measure of DA activity. Hence, an observed BOLD signal change must not necessarily rely on a change in DA transmission, and a change in DA transmission must not necessarily produce a (detectable) BOLD signal change (Brocka et al., 2018). Future research should therefore complement pharmacological fMRI studies with other in vivo techniques that specifically monitor local changes in DA activity, such as molecular imaging with PET and SPECT (single photon emission computed tomography) in humans (Cropley, Fujita, Innis, & Nathan, 2006).

### Conclusion

The present study examined the causal role of DA in human explore/exploit behavior in a pharmacological model-based fMRI approach, using the dopamine precursor L-dopa and the D2 antagonist haloperidol in a placebo-controlled, within-subjects design. First, our cognitive modeling results confirm that humans use both random and uncertainty-based exploration to solve the explore/exploit tradeoff. Notably, we extend previous findings by showing that accounting for choice perseveration improves model fit and interpretability of the parameter capturing uncertainty-based exploration. Our results support the notion that DA is causally involved in the explore/exploit trade-off in humans by regulating the extent to which subjects engage in uncertainty-based exploration. Interestingly, our neuroimaging data do not support the hypothesis that DA controls this trade-off by modulating the neural signatures of exploratory and exploitative decisions per se. In contrast, we provide first evidence that DA modulates tracking of overall uncertainty in a cortical control network comprising the insula and dACC, which might then drive exploration in the face of accumulating uncertainty. Future research should more closely examine the potential role of these regions in driving exploration based on emotional responses to increasing uncertainty, and further investigate how prefrontal and/or striatal DA may be involved in this process.

## Methods

### Participants

In total, 34 healthy male subjects participated in the study (aged 19 to 35 years, M = 26.85, SD = 4.01). Three subjects dropped out of the study due to illness or personal reasons, two after the initial baseline session and one after the first fMRI session. Only males were included, as female sex hormones fluctuate during menstrual cycle which may affect DA signaling (Almey, Milner, & Brake, 2015; Yoest, Quigley, & Becker, 2018). Participants were recruited online and included mainly university students. Inclusion criteria were the following: male, age 18-35 years, normal weight (BMI 18.5-25.0), right-handed, fluent German in speaking and writing, normal or corrected to normal vision, no hearing impairments, no major past or present psychological, neurological, or physical disorders, non-smoker, no excessive consumption of alcohol (<10 glasses per week), no consumption of illegal drugs or prescription drugs within two months prior to the study, no irreversibly attached metal in or on the body, no claustrophobia (the latter two due to the fMRI measurement). Before participating in the study, all subjects provided informed written consent and had to pass a medical check by a physician including an electrocardiogram (ECG) and an interview about their medical history and present health status. Participants received a fixed amount (270€) plus a variable bonus depending on task performance (30-50#x20AC;). The study procedures were approved by the local ethics committee (Hamburg Medical Council).

### General procedure

We employed a double-blind, placebo-controlled, counterbalanced, within-subjects design. Each subject was tested in four separate sessions: one baseline session and three fMRI sessions. The baseline screening was scheduled five to six days prior to the first fMRI session.

### Baseline screening

The baseline screening started with spontaneous eye blink rate assessment, followed by a computerized testing of working memory capacity comprising four working memory tasks, two tasks testing delay and probability discounting behavior (not reported here), and it ended with a psychological questionnaire battery. Participants were encouraged to take small breaks in between the tasks to aid concentration. Blink rate was measured via electromyography for 5 min under resting conditions via three Ag/AgCl electrodes (Blumenthal et al., 2005) and an MP100 hardware system running under AcqKnowledge (version 3.9.1; Biopac Systems, Goleta, CA). Working memory capacity was based on the following tasks: Rotation Span (Foster et al., 2015), Operation Span (Foster et al., 2015), Listening Span (van den Noort, Bosch, Haverkort, and Hugdahl, 2008; based on the English version by Daneman and Carpenter, 1980), and Digit Span (Wechsler Adult Intelligence Scale: WAIS-IV; Wechsler, 2008). All tasks were implemented using the software MATLAB (R2014b; MathWorks, Natick, MA) with the Psychophysics Toolbox extensions (version 3.0.12; Brainard, 1997; Kleiner et al., 2007).

At the end of the baseline screening, subjects completed a computer-based questionnaire battery assessing demographics, personality traits, addictive behavior, and various symptoms of psychopathology. Most of the questionnaires were assessed for a separate study and are of no further importance here. Only the Symptom Checklist-90-Revised (SCL-90-R; Derogatis, 1992; German version by Franke, 1995) was used to exclude subjects with psychiatric symptoms. A cut-off was calculated for each subject and transformed into T values based on a German norm sample of male students (see SCL-90-R manual by Franke, 2000, p. 310-329). As instructed in this manual, the screening cut-off was set to T_GSI_≥63 or T≥63 for at least two of the nine subscales, which was reached by none of the participants. Further, the Edinburgh Handedness Inventory (Oldfield, 1971) was used to ensure that all participants were right-handed.

### Scanning procedure

In the three fMRI sessions, each participant performed two tasks inside the MRI scanner under the three different drug conditions. The procedure for each scanning session was as follows: Upon arrival (2.5 hours before testing in the MRI scanner), participants received a first pill containing either 2 mg haloperidol or placebo (maize starch). Two hours later, subjects received a second pill containing either Madopar (150 mg L-dopa + 37.5 mg benserazide) or placebo. Over the course of the study, each subject received one dose of Madopar in one session, one dose of haloperidol in another session, and two placebo pills in the remaining session (counterbalanced). Half an hours later, subjects first performed the restless four-armed bandit task, followed by an additional short reinforcement learning task (not reported here) both inside the MRI scanner. Both tasks were trained on a practice version outside the scanner a priori. On the first fMRI session, a structural MR image (T1) was additionally obtained. Each fMRI session ended with a post-fMRI testing outside the scanner, to assess several control variables (see *SI*). Throughout each fMRI session, further control variables were assessed at different time points, including physical wellbeing parameters and mood (see *SI*). Subjects were not allowed to eat or drink anything but water throughout the fMRI session but were offered a small snack (cereal bar) after testing in the fMRI scanner to aid concentration for the post-fMRI testing.

### Restless four-armed bandit task

The restless four-armed bandit task was adapted from Daw et al. (2006). The task included 300 trials, which were separated by short breaks into four blocks à 75 trials. Each trial started with the presentation of four different colored squares (“bandits”) representing four choice options (Figure 9a). The squares were displayed on a screen that was reflected in a head coil mirror inside the fMRI scanner. Participants selected one option using a button box held in their right hand. Subjects had a maximum of 1.5 s to indicate their choice. If no button was pressed during that time, a large red X was displayed 4.2 s in the center of the screen indicating a missed trial with no points earned. If subjects pressed a button before the response deadline the selected bandit was highlighted with a black frame. After a waiting period of 3 s during which three black dots were shown within the chosen bandit, the number of points earned in this trial was displayed within the chosen bandit for 1 s. Subsequently, the bandits disappeared, and a fixation cross remained on screen until the trial ended 6 s after trial onset. This was followed by a jittered inter-trial interval (poisson distribution, mean: 2 s (0-5 s)). At the end of the task, the sum of points earned as well as the monetary payout resulting from these points were displayed on screen. Participants were told in advance that 5 % of all points earned would be paid out after the experiment (5 cents per 100 points). The mean payoffs of the four bandits drifted randomly across trials according to a decaying Gaussian random walk. We used the three instantiations from Daw et al. (2006) for the three fMRI sessions of the current study. One of these instantiations is shown in Figure 9b. The order of these three instantiations across fMRI sessions was the same for all subjects, thereby unconfounded with the drug order, which was counterbalanced across subjects. The task was implemented using the software MATLAB (R2014b; MathWorks, Natick, MA) with the Psychophysics Toolbox extensions (version 3.0.12; Brainard, 1997; Kleiner et al., 2007).

**Figure 9.**
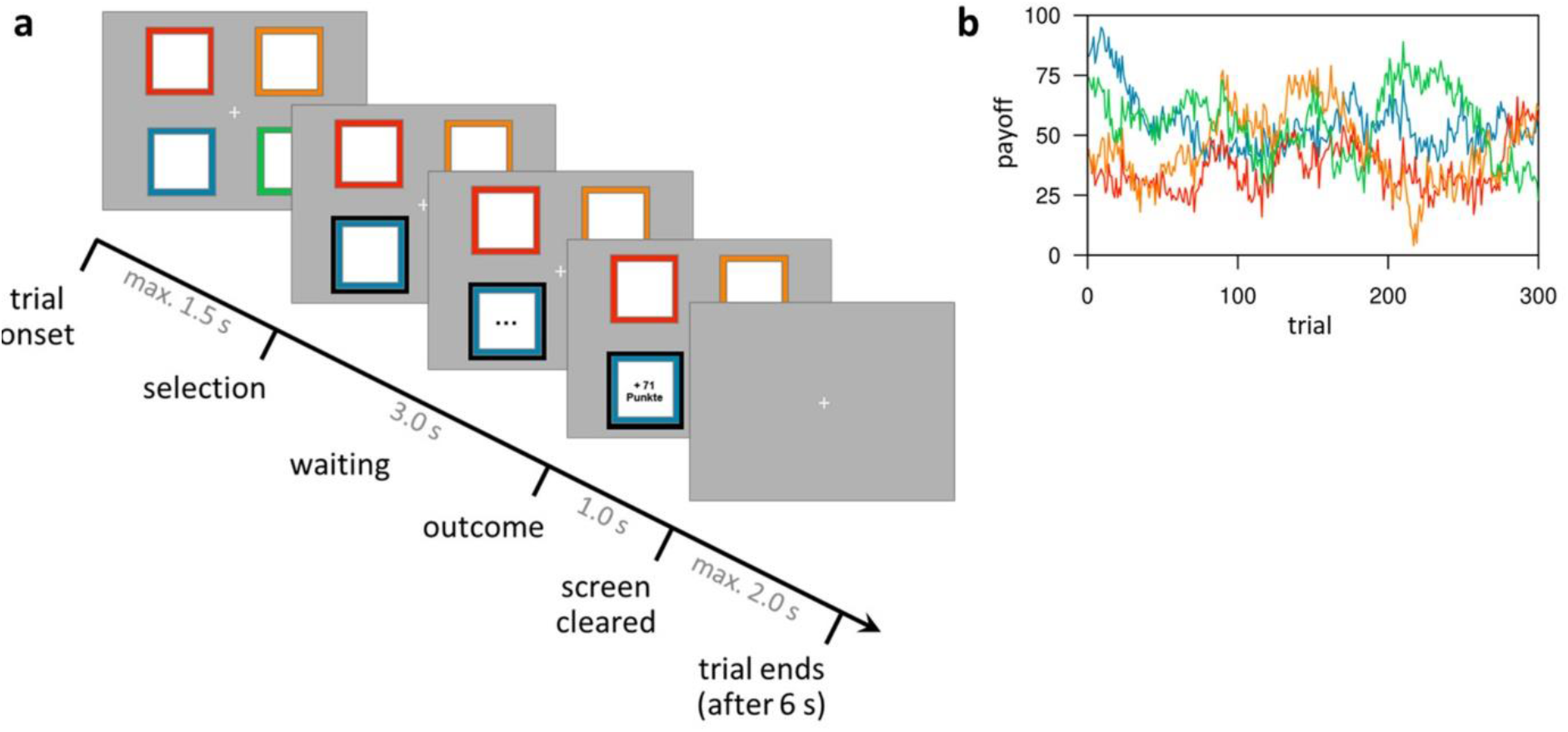
Task design of the restless four-armed bandit task (Daw et al., 2006). (a) Illustration of the timeline within a trial. At trial onset, four colored squares (bandits) are presented. The subject selects one bandit within 1.5 s, which is then highlighted and, after a waiting period of 3 s, reveals its payoff for 1 s. After that, the screen is cleared and the next trial starts after a fixed trial length of 6 s plus a variable intertrial interval (not shown) with a mean of 2 s. (b) Example of the underlying reward structure. Each colored line shows the payoffs of one bandit (mean payoff plus Gaussian noise) that would be received by choosing that bandit on each trial.

### Computational modeling of explore/exploit behavior

Choice behavior in the four-armed bandit task was modeled using six different computational models of explore/exploit choice behavior (see table 1). The best fitting model (in terms of predictive accuracy) was selected for subsequent analyses of behavioral and fMRI data, and pharmacological intervention effects. Each computational model was composed of two components: First, a learning rule (Delta rule, Bayesian learner) described how participants generate and update subjective reward value estimates for each choice option (bandit) based on previous choices and obtained rewards. Second, a choice rule (softmax, softmax + exploration bonus, softmax + exploration bonus + perseveration bonus) modeled how these learned values influence choices. By combining two different learning rules with three different choice rules, a total of six models entered for model comparison.

For the sake of brevity, here we only outline the architecture of the Bayesian learner models (see *SI* for the models implementing the Delta rule), which consistently outperformed the Delta rule models (Daw et al., 2006). This model implements the Kalman filter (Kalman, 1960; Kalman & Bucy, 1961; Anderson & Moore, 1979) as the Bayesian mean-tracking rule for the reward-generating diffusion process in the bandit task. The model assumes that subjects form an internal representation of the true underlying reward structure of the task. The payoff of in trial *t* of bandit *i* followed a decaying Gaussian random walk with mean payoff *μ_i,t_* and variance 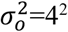 (observation variance). From one trial to the next, the mean payoffs changed according to: *μ*_*i,t*+1_ = *λμ*_*i,t*_ + (1 − *λ*)*ϑ* + *v_t_* with parameters *Λ*=0.9836 (decay parameter), *ϑ*=50 (decay center), and diffusion noise *v_t_* drawn independently in each trial from a Gaussian distribution with zero mean and 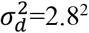(diffusion variance). In the model, subjects’ estimates of these parameters are denoted accordingly as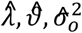 and 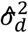. According to the model, participants update their reward expectations of the chosen bandit according to Bayes’ theorem. They start each trial with a prior belief about each bandit’s mean payoff, that is normally distributed with mean 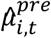 and variance 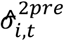 for bandit *i* on trial *t*. For the chosen bandit, this prior distribution is updated by the reward observation *r_t_*, resulting in a posterior distribution with mean 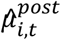 and variance 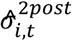 according to:

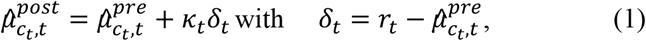

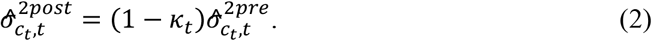

Here *κ* denotes the Kalman gain that is computed for each trial *t* as:

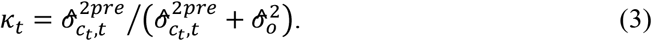

The Kalman gain determines the fraction of the prediction error that is used for updating. In contrast to the learning rate (Delta rule), the Kalman gain varies from trial to trial depending on the current variance of the expected reward’s prior distribution 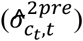 and the estimated observation variance 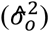. The observation variance indicates how much the actual rewards vary around the (to be estimated) mean reward of a bandit and therefore reflects how reliable each trial’s reward observation (each new data point) is for estimating the true underlying mean. If the prior variance is large compared to the estimated observation variance, i.e. if a subject’s reward prediction is very uncertain while the reward observation is very reliable, the Kalman gain approaches 1 and a large fraction of the prediction error is used for updating. If, in contrast, the prior variance is very small compared to the estimated observation variance, i.e. if a subject’s reward estimation is very reliable while reward observations are very noisy, then the Kalman gain approaches 0 and only a small fraction of the prediction error is used for updating. Similar to the Delta rule, the expected rewards (prior mean and variance) of all unchosen bandits are not updated *within* a trial, i.e. their posteriors equal the prior for that trial. However, prior distributions of all four bandits are updated *between* trials based on the subject’s belief about the underlying Gaussian random walk by:

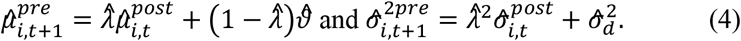

The trial-by-trial updating process was initialized for all bandits with the same prior distribution 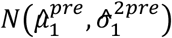, with 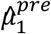 and 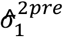 as additional free parameters of the model.

The three choice rules were based on the softmax function (McFadden, 1973; Sutton & Barto 1998). The first implementation utilized the softmax (SM) with the inverse temperature parameter β modeling inherent choice randomness (random exploration). The second choice rule (SM+E) modeled uncertainty-based exploration in addition and the third choice rule further modeled choice perseveration (SM+E+P), both via a bonus that was added to the expected value. The resulting probabilities *P_i,t_* to choose bandit *i* on trial *t* were then:

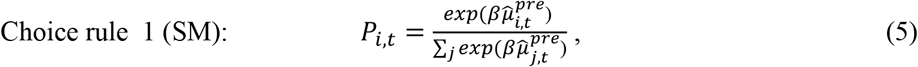

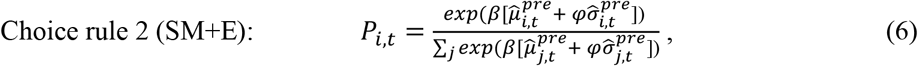

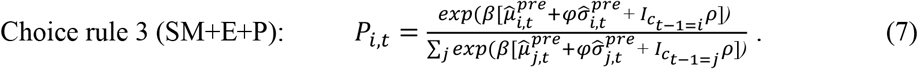

Choice rule 2 is the “softmax (random exploration) with exploration bonus”, as used in Daw et al. (2006). Here, *φ* denotes the exploration bonus parameter, which reflects the degree to which choices are influenced by the uncertainty associated with each bandit.

Choice rule 3 is a novel extension of this model called “softmax with exploration and perseveration bonus”. It includes an extra perseveration bonus, which is a constant value (free parameter) only added to the expected value of the bandit chosen in the previous trial. Here, *ρ* denotes the perseveration bonus parameter and *I* an indicator function that equals 1 for the bandit that was chosen in the previous trial (indexed by *c*_*t*−1_) and 0 for all other bandits.

As mentioned above, all three choice rules were also implemented within a simple Delta rule learning scheme (see *SI*). Taken together, by combing each learning rule with each choice rule, six cognitive models entered model comparison. The parameters for each model are summarized in Table 1.

Posterior parameter distributions were estimated for each subject and drug condition using hierarchical Bayesian modeling within Stan (version 2.17.0; Stan Development Team, 2017b), operating under the general statistical package R (version 3.4.3; R Core Team, 2017). Stan is based on Hamiltonian Monte Carlo sampling (Girolami & Calderhead, 2011) for approximation. Sampling was performed with four chains, each chain running for 1000 iterations without thinning after a warmup period of 1000 iterations. The prior for each group-level mean was uniformly distributed within the limits as given in Figure 11. For each group-level standard deviation, a half-Cauchy distribution with location parameter 0 and scale parameter 1 was used as a weakly informative prior (Gelman, 2006). Priors for all subject-level parameters were normally distributed with a parameter-specific mean and standard deviation (denoted by *M^x^* and Λ^*x*^ for any parameter *x*). Group-level posterior distributions of the three parameters (*β, φ, ρ*; mean and standard deviation) were estimated separately for each drug condition, which allowed the comparison of subject-level as well as group-level parameters between drugs (for details on the fixed parameters see *SI*).

**Table 1.** Free and fixed parameters of all six computational models.

|  | Delta rule |  | Bayes learner rule |  |
| --- | --- | --- | --- | --- |
| Choice rule 1 | $\alpha, \beta$ | *fixed: $v_1$* | $\beta$ | *fixed: $\hat{\lambda}, \hat{\vartheta}, \sigma_o^2, \sigma_d^2, \mu_1^{pre}, \sigma_1^{pre}$* |
| Choice rule 2 | $\alpha, \beta, \varphi$ | *fixed: $v_1$* | $\beta, \varphi$ | *fixed: $\hat{\lambda}, \hat{\vartheta}, \sigma_o^2, \sigma_d^2, \mu_1^{pre}, \sigma_1^{pre}$* |
| Choice rule 3 | $\alpha, \beta, \varphi, \rho$ | *fixed: $v_1$* | $\beta, \varphi, \rho$ | *fixed: $\hat{\lambda}, \hat{\vartheta}, \sigma_o^2, \sigma_d^2, \mu_1^{pre}, \sigma_1^{pre}$* |

*Note:* Free parameters are only listed for the subject-level. Hierarchical models contained for each free subject-level parameter $x$ two additional free parameters ( $M^x, \Lambda^x$ ) on the group-level (Figure 10). Choice rule 1: softmax; Choice rule 2: softmax with exploration bonus; Choice rule 3: softmax with exploration bonus and perseveration bonus; $\alpha$ : learning rate; $\beta$ : softmax parameter; $\varphi$ : exploration bonus parameter; $\rho$ : perseveration bonus parameter; $v_1$ : initial expected reward values for all bandits; $\hat{\lambda}$ : decay parameter; $\hat{\vartheta}$ : decay center; $\sigma_o^2$ : observation variance; $\sigma_d^2$ : diffusion variance; $\mu_1^{pre}$ : initial mean of prior expected rewards for all bandits; $\sigma_1^{pre}$ : initial standard deviation of prior expected rewards for all bandits

### Model comparison

Following parameter estimation, the six cognitive models were compared in terms of predictive accuracy using a Bayesian leave-one-out (LOO) cross-validation approach (Vehtari, Gelman, & Gabry, 2017). LOO cross-validation computes pointwise out-of-sample predictive accuracy by repeatedly taking one data set (“testing set”) out of the sample, refitting the model to the reduced data (“training set”), and then measuring how accurately the refitted model predicts the data of the testing set. A testing set was defined as the data of one subject under one drug condition, compounded over all trials. Model comparison was performed using the data sets from all 31 participants once combined over all drug conditions (yielding 93 data sets) and once separately for each drug condition (each with 31 data sets). To reduce computational burden, the R package loo (Vehtari et al., 2017) was used, which applies Pareto-smoothed importance sampling to calculate LOO estimates as a close approximation. LOO estimates were calculated for each model fit based on its Stan output, using the log likelihood function evaluated at the sampled posterior parameter values. The log likelihood for each subject was calculated as the logarithmized product of choice probabilities (*P*) of the chosen bandits (indexed by *c_t_*) compounded over trials: log(⊓_*t*_*P_c_t_,t_*). Please note that since cross-validation measures like LOO are not biased in favor of more complex models (like ordinary goodness-of-fit measures), no penalty term is needed here to compensate for model complexity in order to prevent overfitting. Based on the results of the model comparison (Figure 1), the cognitive model with the highest predictive accuracy (Bayes-learner with exploration and perseveration bonus (choice rule 3)) was then selected for further data analysis.

**Figure 10.**
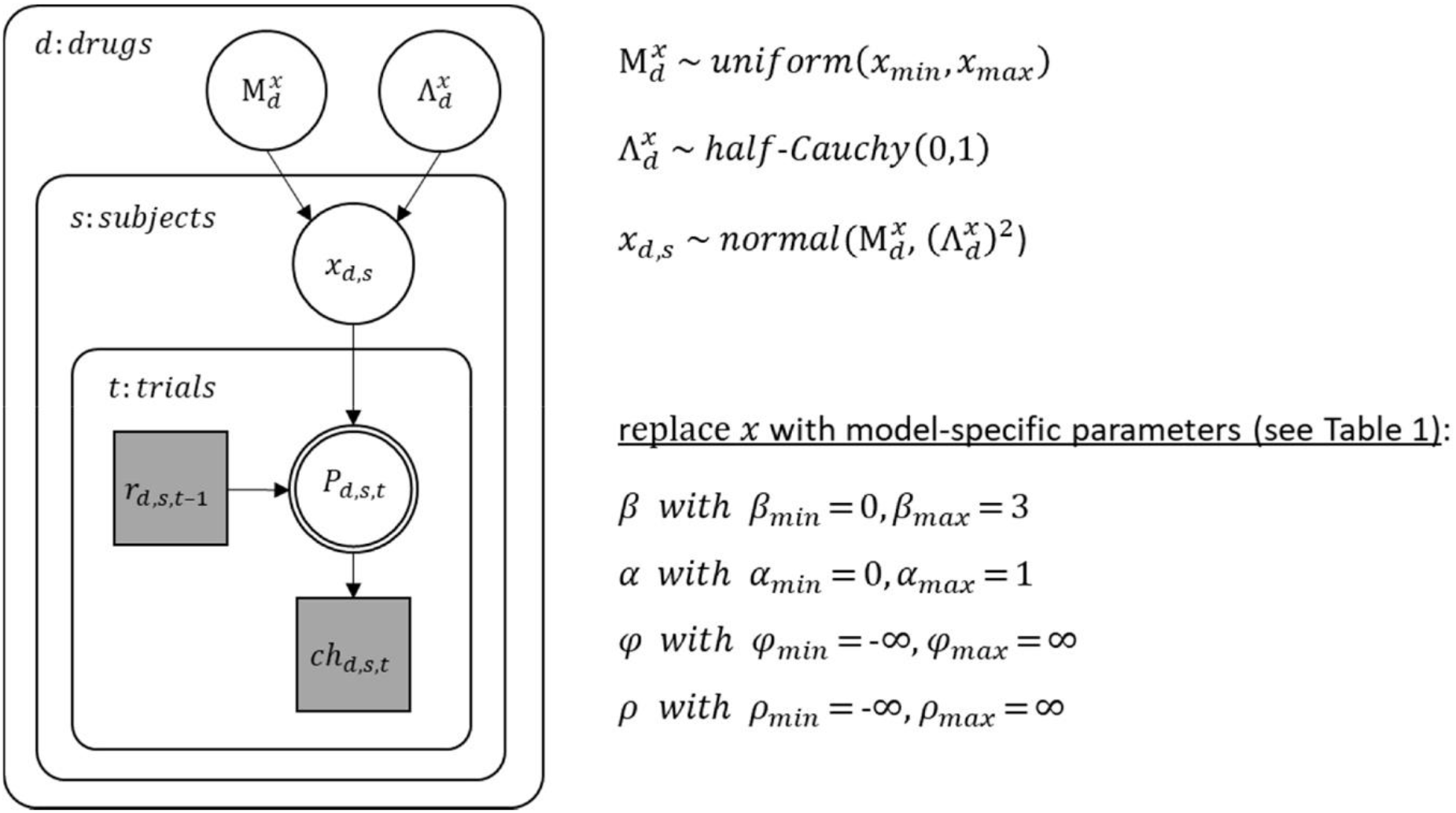
Graphical description of the hierarchical Bayesian modeling scheme. In this graphical scheme, nodes represent variables of interest (squares: discrete variables; circles: continuous variables) and arrows indicate dependencies between these variables. Shaded nodes represent observed variables, here rewards (*r*) and choices (*ch*) for each trial (*t*), subject (*s*), and drug condition (*d*). For each subject and drug condition, the observed rewards until trial t-1 determine (deterministically) choice probabilities (*P*) on trial *t*, which in turn determine (stochastically) the choice on that trial. The exact dependencies between previous rewards and choice probabilities are specified by the different cognitive models and their model parameters (*x*). Note that the double-bordered node indicates that the choice probability is fully determined by its parent nodes, i.e. the reward history and the model parameters. As the model parameters differ between all applied cognitive models, they are indicated here by an *x* as a placeholder for one or more model parameter(s). Still, the general modeling scheme was the same for all models: Model parameters were estimated for each subject and drug condition and were assumed to be drawn from a group-level normal distribution with mean *M^x^* and standard deviation Λ^*x*^ for any parameter *x*. Note that group-level parameters were estimated separately for each drug condition. Each group-level mean (*M^x^*) was assigned a non-informative (uniform) prior between the limits *x_min_* and *x_max_* as listed above. Each group-level standard deviation (Λ^*x*^) was assigned a half Cauchy distributed prior with location parameter 0 and scale 1. Subject-level parameters included *α, β, φ* and *ρ*, depending on the cognitive model (see Table 1).

### FMRI data acquisition

Functional imaging data were acquired on a Siemens Trio 3T scanner (Erlangen, Germany) equipped with a 32 channel head-coil. For each subject and drug condition, four blocks à 75 trials were recorded for the bandit task. The first five scans of each block served as dummy scans to allow for magnetic field saturation and were discarded. Functional volumes were recorded using a T2*-weighted EPI sequence. Each volume consisted of 40 slices with 2mm isotropic voxels and 1mm gap, acquired with a repetition time of 2470ms, an echo time of 26 ms, and a flip angle of 80°. In addition, a high-resolution structural image was acquired for each subject at the end of the first fMRI session, using a T1-weighted magnetization prepared rapid gradient echo (MPRAGE) sequence with 1mm isotropic voxels and 240 slices. The experimental task was projected onto a mirror attached to the head coil and participants responded by using a button box with four buttons held in the right hand.

### FMRI data analysis

#### Preprocessing

Preprocessing and statistical analysis of fMRI data was performed using SPM12 (Wellcome Department of Imaging Neuroscience, London, UK). The preprocessing included four steps: (1) realignment and unwarping to the first image of the placebo session; (2) slice time correction to the onset of the middle slice; (3) spatial normalization to Montreal Neurological Institute (MNI) space utilizing the DARTEL approach (Ashburner, 2007) with a resampling of functional images to 1.5 mm isotropic resolution; (4) spatial smoothing using a Gaussian kernel of 6mm full-width at half-maximum (FWHM).

#### First-level analysis

For the first-level analysis of fMRI data, a general linear model (GLM) was created for each subject and drug condition, concatenated over all four blocks of the bandit task. For each trial in which a bandit was chosen, two different time points were included in the model: trial onset and outcome presentation. GLM regressors for these time points were created by convolving these event onsets (stick function of zero duration) with the canonical hemodynamic response function (HRF). Parametric modulators for both onset regressors were included in the model: (1) the type of each choice (1=explore, 0=exploit, see Daw et al., 2006) as a parametric modulator for the trial onset regressor; (2) the reward prediction error and the outcome value as separate parametric modulators for the outcome regressor. For trials in which no bandit was chosen, the model contained an additional error regressor. Four sessions constants (not convolved with the HRF) were included in the model. Low-frequency noise was removed by employing a temporal high-pass filter with a cut-off frequency of 1/128Hz, and a first order autoregressive model AR(1) was used to remove serial correlations. Regressor-specific contrast images were created for each subject and drug condition for the five regressors of interest: trial onset, outcome onset, choice type, prediction error, and outcome value.

In addition to the main GLM, two alternative GLMs were created. Both alternative GLMs only differed from the main GLM with respect to the regressors modeled at trial onset, while the remaining regressors were the same. Whereas the main GLM included one trial onset regressor with one parametrical modulator (explore/exploit), the second GLM included instead three trial onset regressors: one for uncertainty-based explorations (uncertainty-based), one for random explorations (random), and one for exploitations (exploit). These three choice types were defined according to the trinary classification scheme as described in the results section. The third GLM included one trial onset regressor with two parametric modulators: the expected value 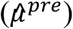 and uncertainty 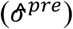 of the chosen option (in that order), both derived from the cognitive model as described in the results section.

#### Second-level analysis

Utilizing a second-level random effects analysis approach, the subject- and drug-specific contrast images for each first-level regressor were submitted to a flexible factorial model in SPM12, including the factors drug (3 levels, within-subject), subject (31 levels), and a constant. For each contrast-specific second-level analysis, a t-contrast image was created that tested for the main effect of that specific contrast over all subjects and drug conditions, calculated by weighting each drug level by one and each subject level by 3/31 (Gläscher & Gitelman, 2008). For the choice type regressor (explore=1, exploit=0), t-contrast were computed twice, once with positive and once with negative weights to create t-contrast images for both comparisons explore>exploit and exploit > explore.

For the second GLM, the second-level random effects analysis included the t-contrasts uncertainty-based > exploit, random> exploit, uncertainty-based > random, and random>uncertainty-based. For the third GLM, t-contrasts for both parameteric modulators, i.e. expected value and uncertainty, were included in the second-level random effects analysis.

To test for DA drug effects across subjects, an F-contrast image was created for each contrast-specific second-level analysis with the weights [1 −1 0; 0 1 −1] over the three drug levels [P D H] and zero weights for all 31 subject levels (Henson & Penny, 2005).

In addition, a second-level regression analysis was conducted for each drug pair to test whether DA drug effects on exploration-specific brain activity were linearly predicted by DA drug effects on exploratory behavior. For this, the subject- and drug-specific contrast images for explore vs. exploit were used to calculate the difference image of this contrast for a given drug pair (P-D, P-H, or D-H) for each subject. These difference images entered a second-level regression analysis, including the subject specific drug differences of the exploration bonus parameter *φ* posterior medians for the same drug pair as explanatory variable. The same kind of regression analysis was also performed for the contrasts uncertainty-based vs. exploit and random vs. exploit of the second GLM.

Post-hoc, a fourth first-level GLM was created for an additional exploratory analysis. This fourth GLM differed from the main GLM only with respect to the parametric modulator of the trial onset regressor, replacing the binary variable choice type (explore/exploit) by a continuous model-based variable termed overall uncertainty 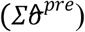, which is the summed uncertainty 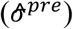 over all four bandits. The contrast images for the overall uncertainty regressor were then used in a second-level random effects analysis to test for drug differences in the parametric effects of this regressor across subjects. Since this post-hoc analysis specifically focused on a comparison of the placebo and L-dopa condition (based on the behavioral findings, see 5.2), the second-level model only included these two drug conditions. Based on this model, different t-contrast images were created to test for the parametric effects of this regressor in the placebo condition alone, and for its differential parametric effects between both drug conditions (placebo>L-dopa, L-dopa>placebo).

For completeness, also a second-level analysis with all three drug conditions was performed to test for the remaining pairwise drug effects accordingly (placebo>haloperidol, haloperidol >placebo, L-dopa>haloperidol, haloperidol> placebo). Finally, also a second-level regression analysis was performed for this regressor.

All fMRI results are reported at a threshold of p<.05, FWE-corrected for the whole brain volume, unless stated otherwise. In addition, results of the second-level ANOVA and regression analysis for the first and second GLM (i.e. exploration-specific contrasts) were also analyzed using small volume FWE correction (p<.05) for seven regions that have previously been associated with exploratory choices: the left/right FPC and left/right IPS (Daw et al., 2006), as well as the dACC and left/right AI (Blanchard & Gershman, 2018). Regions used for small volume correction were defined by a 10 mm radius sphere around the respective peak voxel reported by the previous studies (see *SI*). For display purposes, an uncorrected threshold of p<.001 was used (unless stated otherwise), and activation maps were overlaid on the mean structural scan of all participants.

## Acknowledgements

This work was funded by Deutsche Forschungsgemeinschaft (grant PE1627/5-1 to J.P.). Large parts of this publication are based on the dissertation “Dopaminergic modulation of the explore/exploit trade-off in human decision making” by K.C. (Chakroun, Karima: Dopaminergic modulation of the explore/exploit trade-off in human decision making [online]; Hamburg, Univ., Diss., 2019; URL: http://ediss.sub.uni-hamburg.de/volltexte/2019/9835/)

