## supplementary information for "Dopaminergic modulation of the exploration/exploitation trade-off in human decision-making"

#### Results

##### Correspondence between model parameters and fraction of random exploration, uncertainty-based exploration and exploitation trials

To verify the correspondence between the trinary trial classification (random exploration, uncertainty-based exploration, exploitation) and the computational modeling, we used Pearson correlations to test for associations between the model parameters (subject-level medians of β, $\varphi$, ρ) and the percentage of the three choice types. To increase the sample size for this correlation, we combined data from the placebo condition (n=31) with data from a prior pilot study using the same task (n=16). With the original binary classification (Daw et al., 2006), the percentage of exploration trials per subject was negatively correlated with the random exploration parameter β, but positively correlated with the directed exploration parameter $\varphi$ (Table S1). However, with the trinary classification, only the percentage of random explorations correlated with β, whereas directed explorations were significantly associated with $\varphi$ (Table S1), indicating that both parameters indeed reflect different types of exploration.

**Table S1.** Pearson correlation coefficients between the percentage of exploratory trials per subject and the subject-level parameter medians of the winning model.

| **% explorations** | **β** | **ϕ** | **ρ** |
|---|---|---|---|
| **overall** | **-.65*****| **.30*** | -.18 |
| **random** | **-.68***** | -.09 | -.22 |
| **directed** | -.28 | **.64***** | -.09 |

Note that overall explorations were defined according to the binary choice classification, while directed and random explorations were defined according to the trinary choice classification. β: RE parameter; $\varphi$: DE parameter; ρ: CP parameter. *p<.05. ***p<.001.

##### Drug-effects on the fraction of random exploration, uncertainty-based exploration and exploitation trials

We complemented the analyses of drug effects on model parameters (β, $\varphi$, ρ), with an analysis of drug effects on the percentage of exploitation and exploration trials (overall, random and uncertainty-based) per subject. For this, a rmANOVA with within factor drug was performed for each of the following four dependent variables: (a) the percentage of overall exploration trials, (b) the percentage of random exploration trials, (c) the percentage of uncertainty-based exploration trials, and (d) the percentage of exploitation trials. In accordance with the previous analysis, we observed a significant drug effect only for the percentage of uncertainty-based explorations ($F_{2,60}$=7.15, p=.002; Figure S1), but not for the percentage of random explorations ($F_{2,60}$=0.53, p=.592), overall explorations (F2,60=0.97, p=.386), or exploitations (F2,60=1.62, p=.207). Post-hoc, paired t-tests showed a significant reduction in the percentage of uncertainty-based explorations under L-dopa compared to placebo (mean difference P-D=2.82, $t_{30}$=4.69, p<.001) and haloperidol (mean difference H-D =2.42, $t_{30}$=2.76, p=.010), but not between placebo and haloperidol (mean difference P-H=0.39, $t_{30}$=0.43, p=.667). Notably, an exploratory t-test revealed that the percentage of exploitations was marginally increased under L-dopa compared to placebo (mean difference P-D=-2.61, $t_{30}$=-1.92, p=.065).

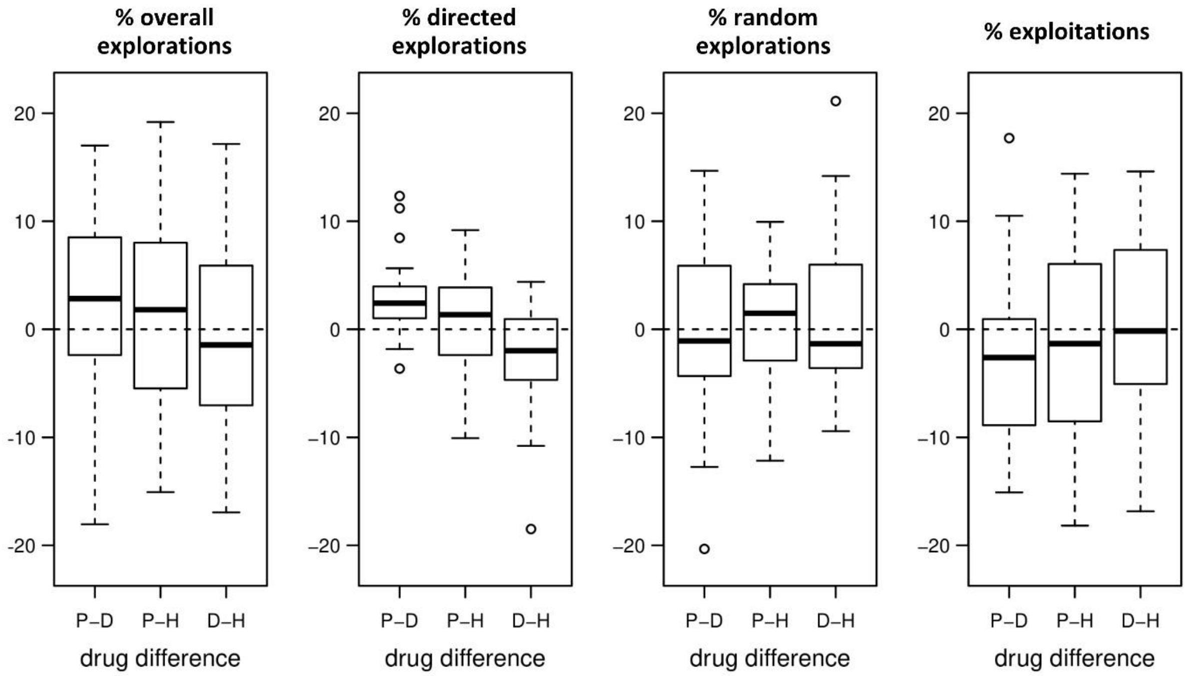


**Figure S1**. Drug effects on the percentage of explorations and exploitations. Shown are the pairwise drug differences for the percentage of overall explorations, directed explorations, random explorations, and exploitations. P: placebo; D: L-dopa; H: haloperidol.

##### Drug-effects on model-free measures of choice behavior

In addition to the model-based choice parameters, also several model-free measures of choice behavior were tested for DA drug effects. These variables included the overall monetary payout (payout), the percentage of choices in which the bandit with the highest actual payoff was selected (%best bandit), the mean rank of all chosen bandits when bandits are ranked by their actual payoff (mean rank), the percentage of switches (%switches), and mean /median reaction times (mean RT, median RT). Yet, the rmANOVA yielded no significant drug effect on any of these four model-free choice variables (payout: $F_{2,60}=0.06$, $p=.943$; %best bandit: $F_{2,60}=0.34$, $p=.711$; mean rank: $F_{2,60}=0.37$, $p=.690$; %switches: $F_{2,60}=1.02$, $p=.366$; mean RT: $F_{2,60}=0.54$, $p=.585$; median RT: $F_{2,60}=0..50$, $p=.611$).

**Drug-effects on control variables**

We checked for drug-related changes on several control variables in addition to the results reported in the main section. The first set of control variables was measured during the post-fMRI testing and included the spontaneous eye blink rate (sEBR), the total scores of the Digit Span Task (forward and backward), and 15 attentional performance measures from the Tests of Attentional Performance (TAP). To test for DA drug effects, a rmANOVA with the factor drug was performed on each of these 18 control variables. None of these 18 variables showed a significant drug effect in the ANOVA (all $p>.05$).

A second set of control variables was measured at different time points throughout each fMRI session and comprised six variables on subjective mood (alertness, contentedness, calmness, pleasure, arousal, and dominance) and four variables on physical wellbeing (pulse, systolic and diastolic blood pressure, and the side effects sum score). To test for drug effects on these variables, their scores at three different time points after drug administration (t1, t2, t3) were subtracted by their baseline score before drug administration (t0) for each drug condition. A rmANOVA with the factor drug on each of these difference scores (t1-t0, t2-t0, t3-t0) showed no significant drug effect for any of the physical wellbeing parameters (all $p>.05$). However, a significant drug effect ($F_{2,60}=4.46$, $p=.016$) was found for one of the difference scores (t3-t0) on the subscale "calmness" of the VAS (Bond & Lader, 1974). Paired t-tests on this variable revealed a significant increase under haloperidol compared to placebo (mean difference P-H= -0.67, $t_{30}=-2.05$, $p=.049$) and L-dopa (mean difference D-H=-0.85, $t_{30}=-2.99$, $p=.005$), but no significant difference between placebo and L-dopa (mean difference P-D=0.18, $t_{30}=0.61$, $p=.544$). Although, with correction for multiple comparisons (in total n=42 tests), the effects no longer remained significant.

Beyond, to test whether subjects were actually blind to the drug condition, their drug guesses after each session were examined. Over all subjects and drug sessions, 30 of the 93 guesses (32.3%) were correct, which is in line with the number of correct guesses expected by chance (31, i.e. 33.3%). To rule out that some subjects performed above chance (i.e. recognized all drug conditions) and others below, the number of correct guesses per subject was calculated, resulting in the following frequency distribution: 25.8% (0 correct guesses), 51.6% (1 correct guess), 22.6% (2 correct guesses), 0.0% (3 correct guesses). A chi-squared test revealed no significant difference to random guessing (29.6% (0 correct guesses), 44.4% (1 correct guess), 22.2% (2 correct guesses), 3.7% (3 correct guesses); $\chi^2$=1.66, p=.634; with Monte Carlo approximation). Finally, a chi-squared test showed that the frequencies of drug guesses did not significantly differ between the three drug conditions ($\chi^2_4$=0.36, p=.986). Taken together, the results of all three analyses indicate that the observed data are in accordance with random guessing.

##### Drug effects on the posterior distributions of the group-level model parameters

In addition to the results reported in the main results section, we here show the drug-specific posterior distributions of the group-level standard deviations of all free model parameters ($\beta, \varphi, \rho$; Figure S2) and the drug-related differences of the group-level mean and standard deviation for parameter $\varphi$ (Table S2).

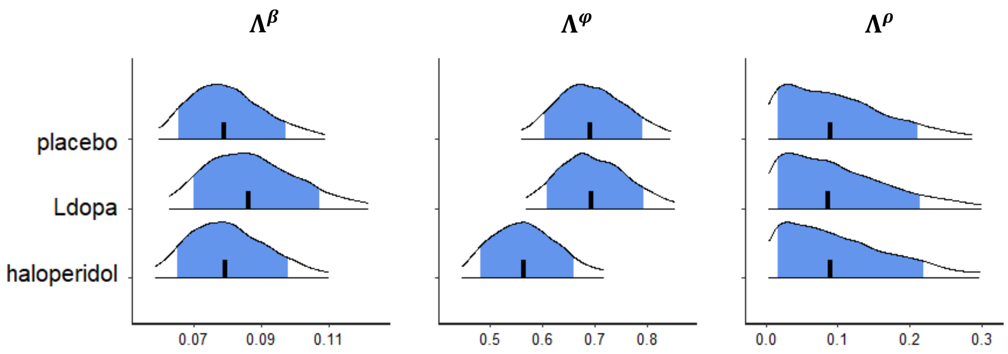


**Figure S2**. Group-level parameter estimates of the winning model. Shown are the posterior distributions of the group-level standard deviation ($\Lambda$) for all choice parameters ($\beta, \varphi, \rho$) of the winning model, separately for each drug condition. For each posterior distribution, the plot shows the median (vertical black line), the 80% central interval (blue area), and the 95% central interval (black contours). $\beta$: softmax parameter; $\varphi$: exploration bonus parameter; $\rho$: perseveration bonus parameter.

**Table S2**. Drug effects on the exploration bonus parameter ($\varphi$) on the group-level.

| | $M^{\varphi}$ | | $\Lambda^{\varphi}$ | |
|---|---|---|---|---|
| | % above 0 | 90% HDI | % above 0 | 90% HDI |
| **placebo - L-dopa** | **97.5** | **[0.05, 0.69]** | 47.5 | [-0.18, 0.16] |
| placebo - haloperidol | 49.3 | [-0.30, 0.27] | 90.0 | [-0.04, 0.29] |
| L-dopa - haloperidol | 1.7 | [-0.70, -0.10] | 90.8 | [-0.02, 0.31] |

Note: Results refer to the posterior drug differences of the group-level mean ($M^{\varphi}$) and standard deviation ($\Lambda^{\varphi}$) for the $\varphi$ parameter of the winning model. For each posterior difference, the table shows the percentage of samples with values above zero (column: % above 0) and the 90% highest density interval (column: 90%HDI).

**No evidence for the inverted-u-shape dopamine hypothesis**

We predicted participants' individual DA baseline proxies (spontaneous eye blink rate (sEBR), working memory capacity (WMC)) to modulate explore/exploit behavior in a quadratic fashion and to predict the strength and direction of drug-related effects in a linear manner. To test the first hypothesis, it was examined whether individual differences in explore/exploit behavior, as assessed by different model-based and model-free choice variables, were predicted by the individual DA baseline, indexed by the sEBR and WMC, according to an inverse quadratic (inverted-U-shape) relationship. To increase sample size for this analysis, data from the placebo condition (n=31) and a pilot study (n=16) were combined, resulting in a sample of 47 subjects. In summary, we found no evidence for an inverted-U-shaped association of choice behavior and DA proxy measures (see Table S3 & Figure S3).

**Table S3**. Test for an inverted-U relationship between choice behavior and DA baseline.

| | **$Loo_{LM}$ - $Loo_{QM}$** | | **ß₂ estimate** | | **ß₂ *p*-value** | |
|---|---|---|---|---|---|---|
| | sEBR | WMC | sEBR | WMC | sEBR | WMC |
| **model-based:** | | | | | | |
| $\beta$ | -0.06 | -0.04 | -2.09e$^{-04}$ | 2.98e$^{-04}$ | .132 | .949 |
| $\varphi$ | -3.60 | -2.57 | -1.13e$^{-03}$ | 1.27e$^{-02}$ | .470 | .809 |
| $\rho$ | -53.09 | -49.68 | 1.69e$^{-03}$ | 1.20e$^{-01}$ | .869 | .726 |
| **model-free:** | | | | | | |
| *payout* | -0.95 | -1.05 | -6.04e$^{-04}$ | 1.37e$^{-02}$ | .582 | .710 |
| *%best bandit* | 198.06 | -245.78 | -2.45e$^{-02}$ | 7.40e$^{-02}$ | .149 | .897 |
| *mean rank* | 0.06 | -0.10 | -5.29e$^{-04}$ | -3.45e$^{-03}$ | .080 | .733 |
| *%switches* | -484.04 | -700.67 | -2.09e$^{-04}$ | 6.58e$^{-01}$ | .222 | .509 |

Note. Choice behavior was assessed by the three choice parameters of the winning (Bayes-SM+E+P) model (upper part) and four model-free choice variables (lower part). Baseline dopamine (DA) function was assessed by the two behavioral DA proxies spontaneous eye blink rate (sEBR) and working memory capacity (WMC). For the latter, the first principal component across three different WMC tasks was used, denoted by $WMC_{PCA}$. The column "$LOO_{LM}$-$LOO_{QM}$" denotes the difference of the squared distances for the linear model (LM) minus the quadratic model (QM) from the leave-one-out (LOO) model comparison. Note that negative values for $LOO_{LM}$ - $LOO_{QM}$ indicate better predictive accuracy of the LM. The columns "$\beta_2$ estimate" and "$\beta_2$ p-value" show for each quadratic model the estimated value and p-value of the $\beta_2$ regression coefficient, respectively. Note that data from a pilot study (n=16) and the placebo condition of the main study were combined for this analysis to increase the sample size to n=47. $\beta$: softmax parameter; $\varphi$: exploration bonus parameter; $\rho$: perseveration bonus parameter

**a model-based choice variables**

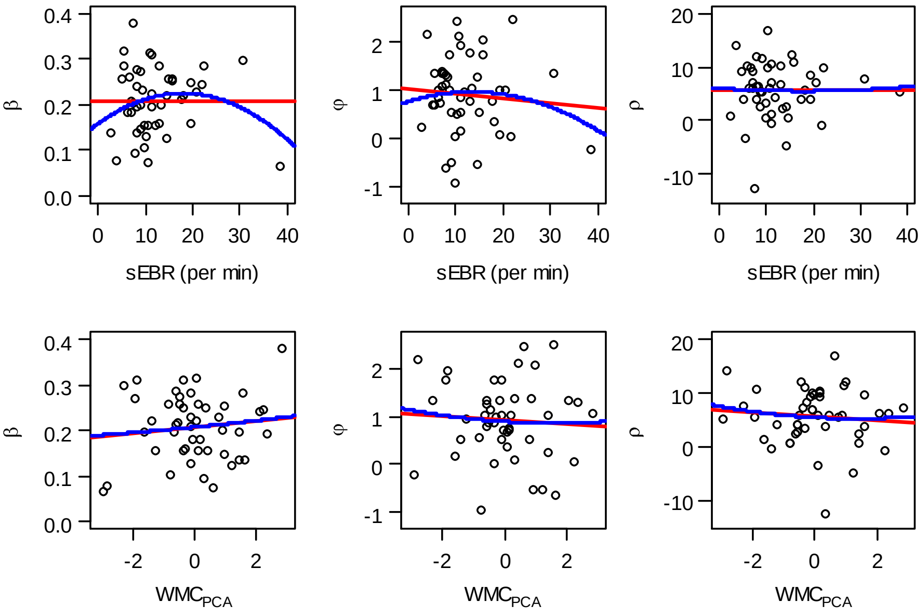


**b model-free choice variables**

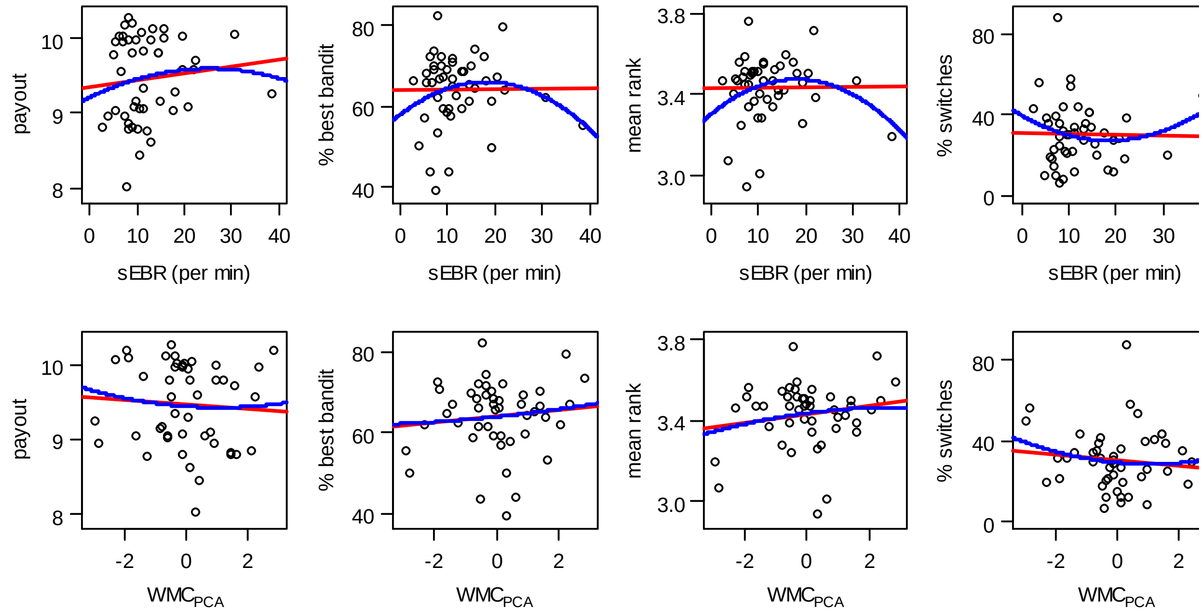


**Figure S3**. Test for an inverted-U relationship between choice behavior and DA baseline. Choice behavior was assessed by (**a**) the posterior medians of the three choice parameters (β, φ, ρ) of the winning (Bayes-SM+E+P) model and (**b**) four model-free choice variables (*payout*, *%best bandit, mean rank, %switches*). DA baseline function was assessed by the two DA proxies spontaneous eyeblink rate (sEBR) and working memory capacity (WMC). For the latter, the first principal component across three different WMC tasks was used, denoted by $WMC_{PCA}$. Each plot shows two regression lines that were fitted to the data, one for the "linear model" (red line) and one for the "quadratic model" (blue line). Note that data from pilot study 2 and the placebo condition of the main study were combined for this analysis to increase the sample size to n=47. $\beta$: softmax parameter; $\varphi$: exploration bonus parameter; $\rho$: perseveration bonus parameter.

Next, we used linear regression models to test the prediction of a linear association between the drug-related differences of subject-level model parameters ($\beta, \varphi, \rho$) and proxies of DA baseline (sEBR, WMC). All regression slopes were not significant different from zero (see Table S4).

**Table S4**. Test for a linear relationship between drug-related effects on model-parameters and DA baseline.

| | **$R^2$** | | **$\beta_1$ estimate** | | **$\beta_1$ *p*-value** | |
|---|---|---|---|---|---|---|
| | sEBR | WMC | sEBR | WMC | sEBR | WMC |
| $\beta$ (P-D) | $2.13e^{-5}$ | $1.52e^{-3}$ | $4.75e^{-05}$ | $2.25e^{-03}$ | .98 | .84 |
| $\varphi$ (P-D) | $1.87e^{-2}$ | $4.25e^{-2}$ | $1.40e^{-02}$ | $-1.19e^{-01}$ | .46 | .27 |
| $\rho$ (P-D) | $5.91e\text{-}^{3}$ | $2.42e\text{-}^{3}$ | $-4.89e^{-03}$ | $-1.76e^{-01}$ | .68 | .79 |
| $\beta$ (P-H) | $2.47e^{-2}$ | $2.93e^{-2}$ | $1.64e^{-03}$ | $1.01e^{-02}$ | .40 | .36 |
| $\varphi$ (P-H) | $9.58e^{-3}$ | $3.36e^{-2}$ | $-1.01e^{-02}$ | $-1.06e^{-01}$ | .60 | .32 |
| $\rho$ (P-H) | $4.61e^{-2}$ | $1.18e^{-2}$ | $-9.01e^{-03}$ | $-2.57e^{-01}$ | .25 | .56 |
| $\beta$ (D-H) | $1.57e^{-3}$ | $7.83e^{-3}$ | $1.57e^{-03}$ | $7.83e^{-03}$ | .43 | .49 |
| $\varphi$ (D-H) | $1.02e^{-3}$ | $2.95e^{-4}$ | $-3.93e^{-03}$ | $1.20e^{-02}$ | .86 | .93 |
| $\rho$ (D-H) | $4.11e\text{-}^{3}$ | $5.69e\text{-}^{4}$ | $-4.07e^{-02}$ | $-8.54e^{-02}$ | .73 | .90 |

Note. Drug-related differences (P: placebo, D: L-dopa, H: haloperidol) of model parameters for all participants (n=31). Baseline dopamine (DA) function was assessed by the two behavioral DA proxies spontaneous eye blink rate (sEBR) and working memory capacity (WMC). For the latter, the first principal component across three different WMC tasks was used, denoted by $WMC_{PCA}$. The column "$R^2$" denotes the $R^2$-values of the linear regressions. The columns "$\beta_1$ estimate" and "β1 p-value" show for each linear model the estimated value and p-value of the $\beta_1$ regression coefficient, respectively. $\beta$: softmax parameter; $\varphi$: exploration bonus parameter; $\rho$: perseveration bonus parameter

#### Distinct brain networks orchestrate exploration and exploitation

The center coordinates of the 10mm spheres used for the regions of interest analysis are reported below (Table S5).

**Table S5.** Regions used for small volume correction.

| **region of small volume correction** | **peak voxel (mm) x** | **y** | **z** | **reference for peak voxel** |
|---|---|---|---|---|
| rFPC (right frontopolar cortex) | 27 | 57 | 6 | Daw et al., 2006 |
| lFPC (left frontopolar cortex) | -27 | 48 | 4 | Daw et al., 2006 |
| rIPS (right intraparietal sulcus) | 39 | -36 | 42 | Daw et al., 2006 |
| lIPS (left intrapareital sulcus) | -29 | -33 | 45 | Daw et al., 2006 |
| rAIns (right anterior insula) | 32 | 22 | -8 | Blanchard & Gershman, 2017 |

| | | | | |
|---|---|---|---|---|
| lAIns (left anterior insula) | −30 | 16 | -8 | Blanchard & Gershman, 2017 |
| dACC (dorsal anterior cingulate cortex) | 8 | 16 | 46 | Blanchard & Gershman, 2017 |

Note: Each small volume correction used a 10-mm-radius sphere around the listed voxel coordinates, which mark brain regions that have previously been associated with exploratory choices.

Table S6 depicts the brain regions that showed higher activity in exploratory compared with exploitative choices as revealed by the first GLM. In Table S7 the opposite contrast is depicted.

**Table S6**. Brain regions showing higher activity for exploratory than exploitative choices (first GLM).

| **Region** | **MNI coordinates** | | | **peak** | **cluster** |
|---|---|---|---|---|---|
| | **x** | **y** | **z** | **z-value** | **extent (k)** |
| R/L intraparietal sulcus, R/L postcentral gyrus, R/L precuneus, L precentral gyrus | -48 | -33 | 52 | 10.45 | 15606 |
| R precentral gyrus | 26 | -8 | 50 | 9.32 | 2297 |
| R/L supplementary motor cortex, R/L dorsal anterior cingulate cortex | 8 | 12 | 45 | 8.47 | 2552 |
| R cerebellum / fusiform gyrus | 18 | -51 | -22 | 8.09 | 2574 |
| R middle frontal gyrus (FPC) | 39 | 34 | 28 | 7.56 | 1291 |
| R cerebellum | 24 | -57 | -54 | 7.35 | 128 |
| L precentral gyrus | -51 | 0 | 34 | 7.31 | 430 |
| L cerebellum, L fusiform gyrus | -40 | -54 | -32 | 7.28 | 1419 |
| L thalamus | -10 | -20 | 6 | 6.96 | 556 |
| R/L calcarine cortex | -8 | -74 | 14 | 6.90 | 1222 |
| R anterior insula | 36 | 20 | 3 | 6.87 | 511 |
| L anterior insula | -36 | 15 | 3 | 6.69 | 557 |
| R precentral gyrus | 51 | 8 | 24 | 6.49 | 434 |
| R thalamus | 10 | -18 | 8 | 6.32 | 331 |
| R cerebellum | 30 | -44 | -48 | 6.24 | 28 |
| L middle frontal gyrus (FPC) | -42 | 27 | 27 | 6.07 | 97 |
| R cerebellum | 14 | -62 | -45 | 5.88 | 61 |
| R pallidum | 15 | 6 | -4 | 5.83 | 25 |
| R calcarine cortex | 9 | -94 | 6 | 5.74 | 104 |
| vermis | 3 | -75 | -34 | 5.70 | 52 |
| R supramarginal gyrus | 51 | -42 | 28 | 5.69 | 46 |
| L middle frontal gyrus (FPC) | -30 | 46 | 15 | 5.67 | 47 |
| L pallidum | -10 | 6 | -4 | 5.64 | 51 |
| R anterior orbital gyrus | 24 | 54 | -9 | 5.60 | 33 |
| L posterior cingulate cortex | -3 | -32 | 26 | 5.51 | 21 |
| L caudate nucleus | -16 | -14 | 18 | 5.33 | 28 |
| R caudate nucleus | 12 | -8 | 16 | 5.24 | 16 |
| L lingual gyrus | -16 | -84 | -12 | 5.21 | 10 |
| R anterior cingulate cortex | 10 | 27 | 21 | 5.13 | 10 |

Note: Thresholded at $p<.05$, FWE-corrected for whole-brain volume, with k≥10 voxels; L: left; R: right.

**Table S7.** Brain regions showing higher activity for exploitative than exploratory choices (first GLM).

| Region | MNI coordinates x | y | z | peak z-value | cluster extent (k) |
|---|---|---|---|---|---|
| L angular gyrus | -42 | -74 | 34 | 8.04 | 2530 |
| L posterior cingulate cortex / precuneus | -6 | -52 | 15 | 7.40 | 1087 |
| R angular gyrus | 52 | -68 | 28 | 7.02 | 185 |
| R postcentral gyrus | 33 | -26 | 54 | 6.80 | 503 |
| R cerebellum | 27 | -78 | -38 | 6.28 | 452 |
| R rostral anterior cingulate cortex | 4 | 18 | -14 | 5.90 | 125 |
| L superior temporal gyrus | -62 | -36 | 3 | 5.89 | 70 |
| L lateral orbital gyrus | -38 | 34 | -14 | 5.81 | 102 |
| R central operculum | 45 | -14 | 20 | 5.73 | 83 |
| L middle temporal gyrus | -62 | -4 | -22 | 5.67 | 193 |
| R/L medial frontal cortex (vmPFC) | -2 | 40 | -10 | 5.67 | 233 |
| L superior frontal gyrus | -10 | 54 | 30 | 5.54 | 20 |
| L superior frontal gyrus | -10 | 51 | 36 | 5.45 | 10 |
| L middle temporal gyrus | -60 | -51 | -2 | 5.38 | 61 |
| R superior temporal gyrus | 52 | -12 | -9 | 5.35 | 25 |
| R middle temporal gyrus | 62 | 4 | -21 | 5.30 | 10 |
| L rostral anterior cingulate cortex | -6 | 46 | 4 | 5.17 | 13 |
| L inferior frontal gyrus | -50 | 27 | 2 | 5.16 | 20 |

Note: Thresholded at $p<.05$, FWE-corrected for whole-brain volume, with k≥10 voxels; L: left; R: right.

#### Similar activation patterns for different exploration types

Overlaying activation maps for uncertainty-based, random, and overall explorations (each contrasted against exploitation) showed a highly similar activation pattern for all three exploration conditions Figure S4) that in each case included the same network (bilateral FPC, IPS, dACC, AI, and thalamus).

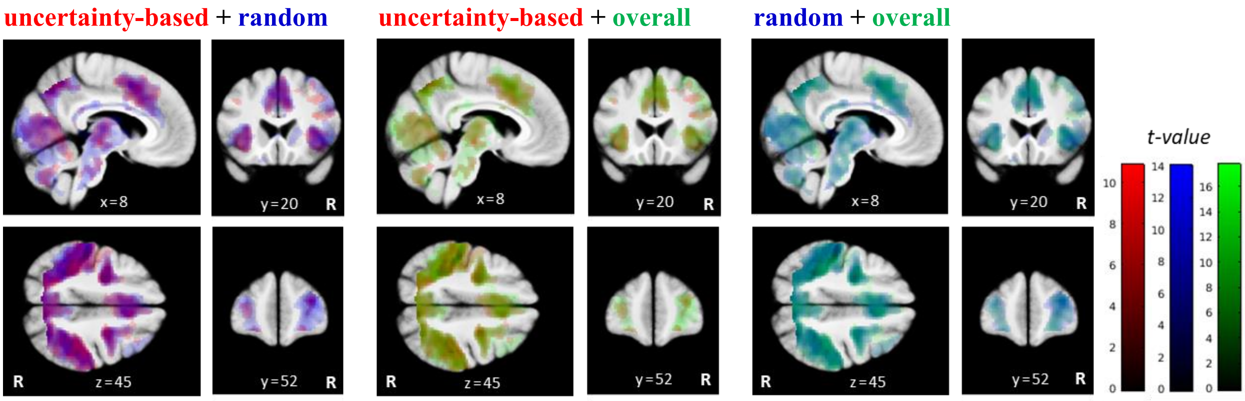


**Figure S4**. Brain activation patterns for different types of explorations**.** Shown are pairwise overlays of the statistical parametric maps for the contrasts explore > exploit ("overall" in green), uncertainty-based > exploit ("uncertainty-based" in red), and random > exploit ("random" in blue) over all drug conditions. While the first contrast is based on a binary choice classification according to which all choices not following the highest expected value are explorations, the other two contrast are based on a trinary choice classification, which further subdivides explorations into choices following the highest exploration bonus (uncertainty-based) and choices not following the highest exploration bonus (random). All activation maps thresholded at p<.05, uncorrected for display purposes. R: right.

**Table S8.** Brain regions in which activity was significantly correlated with the overall uncertainty (fourth GLM), shown for the placebo condition and for pairwise comparison with L-dopa.

| **Region** | **MNI coordinates** | | | **peak** | **cluster** |
|---|---|---|---|---|---|
| | **x** | **y** | **z** | **z-value** | **extent (k)** |
| **placebo** | | | | | |
| L posterior insula | -34 | -20 | 8 | 4.63 | 198 |
| R supplementary motor cortex | 8 | 10 | 52 | 3.98 | 92 |
| R/L dorsal anterior cingulate cortex, L supplementary motor cortex | -3 | 21 | 39 | 3.96 | 176 |
| R anterior insula | 42 | 15 | -6 | 3.46 | 38 |
| R thalamus | 8 | -10 | 2 | 3.41 | 18 |
| **placebo>L-dopa** | | | | | |
| L posterior insula | -34 | -20 | 8 | 5.05[1] | 82 |
| L anterior insula, L frontal operculum | -38 | 6 | 14 | 4.88 | 222 |
| L opercular part of the inferior frontal gyrus | -42 | 9 | 26 | 4.01 | 80 |
| L precentral gyrus | -54 | 3 | 12 | 3.47 | 23 |
| R dorsal anterior cingulate cortex | 4 | 14 | 28 | 3.41 | 32 |
| R precentral gyrus | 39 | -9 | 44 | 3.39 | 16 |
| L dorsal anterior cingulate cortex | -2 | 36 | 33 | 3.32 | 17 |
| **L-dopa>placebo** | | | | | |
| no suprathreshold activation | | | | | |

Note: Thresholded at $p<.001$, uncorrected, with k≥10 voxels; L: left; R: right.

[1] $p=.031$, FWE-corrected for whole-brain volume

### Methods

#### Computational modeling (delta rule)

Choice behavior in the four-armed bandit task was modeled using the combination of two different learning rules (Delta learning rule, Bayesian learner), and three choice rules (softmax, softmax + exploration bonus, softmax + exploration bonus + exploitation bonus) that resulted in six different computational models of explore/exploit choice behavior. Here, the architecture of the delta learning rule (Sutton & Barto, 1998) which is an established temporal difference model of reinforcement learning is outlined.

According to this rule, subjects update the expected reward value ($v$) of a chosen bandit based on their prediction error ($\delta$), i.e. the difference between the actual reward ($r$) and the expected reward for that trial:

$$v_{c_t,t+1} = v_{c_t,t} + \alpha\delta_t \quad \text{with} \quad \delta_t = r_t - v_{c_t,t} \; .$$

Herein, the indices $t$ and $t$+1 denote the current and the next trial, respectively, and $c_t$ the index of the bandit chosen on trial $t$. The parameter $\alpha$ denotes the learning rate, which is a free parameter in this model ranging between 0 and 1. The learning rate determines the fraction of the prediction error that is used for updating. In contrast, the expected rewards of all unchosen bandits are not changed from one trial to the next, i.e. they remain constant until that bandit is chosen again. This trial-by-trial updating was initialized for each bandit with the same expected reward value $v_1$ = 50.

Next, three different choice rules were used to model subjects' choices based on their expected rewards derived from either the Delta rule or the Bayesian learner rule. All choice rules were based on the commonly applied softmax function (McFadden, 1973; Sutton & Barto 1998). The first model was the softmax function (SM) in its basic form without any bonus term. According to this rule, choices are probabilistically based on the relative expected reward values of each choice option. The softmax $\beta$ parameter, also called inverse temperature, reflects the degree of randomness (random exploration) in a subject's decisions. The second choice rule was a modified version of the softmax function called "softmax with exploration bonus" (SM+E), adopted from Daw et al. (2006). This model adds an additional "exploration bonus" to the expected value of each bandit, which increased with the uncertainty of a bandit's outcome. Within the Delta learning rule models, this was accomplished with a simple heuristic adopted from Speekenbrink and Konstantinidis (2015). According to that heuristic, a bandit's uncertainty increases linearly with the number of trials since it was last chosen. This is formalized as $(t - T_i)$, where $T_i$ is the last trial before the current trial $t$ in which bandit $i$ was chosen. The third choice rule was a novel extension of the second choice model called "softmax with exploration and perseveration bonus" (SM+E+P). This version of the softmax rule includes an extra perseveration bonus, in the form of a constant value (free parameter) only added to the expected value of the bandit chosen in the previous trial, but not for all other bandits. In analogy to the Bayseian models (see Methods section of the main article), the three resulting models utilizing the Delta learning rule read as follows:

Choice rule 1 (SM): $$P_{i,t} = \frac{exp(\beta v_{i,t})}{\sum_j exp(\beta v_{j,t})}$$

Choice rule 2 (SM+E): $$P_{i,t} = \frac{exp(\beta[v_{i,t} + \varphi(t-T_i)])}{\sum_j exp(\beta[v_{j,t} + \varphi(t-T_j)])}$$

Choice rule 3 (SM+E+P): $$P_{i,t} = \frac{exp(\beta[v_{i,t} + \varphi(t-T_i) + I_{c_{t-1}=i}\rho])}{\sum_j exp(\beta[v_{i,t} + \varphi(t-T_i) + I_{c_{t-1}=j}\rho])} \; .$$

Herein, $P_{i,t}$ denotes the probability to choose bandit $i$ on trial $t$, and $\sum_j$ indicates a summation over all four bandits; $\varphi$ denotes the exploration bonus parameter, which reflects the degree to which choices are influenced by the uncertainty associated with each bandit and $\rho$ denotes the perseveration bonus parameter and $I$ an indicator function that equals 1 for the bandit that was chosen in the previous trial (indexed by $c_{t-1}$) and 0 for all other bandits.

**Fixed parameters (Bayesian learner)**

For all Bayes learner models, the six parameters specifying subjects' estimation of the Gaussian random walk ($\hat{\lambda}, \hat{\vartheta}, \hat{\sigma}_o^2, \hat{\sigma}_d^2, \hat{\mu}_1^{pre}, \hat{\sigma}_1^{pre}$) ("random walk parameters") were fixed to constrain parameter space for these models, which largely facilitated estimation of the remaining choice parameters. Also, some of the Bayes learner models did actually not converge with free random walk parameters, thus fixing these parameters was necessary in order to include all six cognitive models in the model comparison. In detail, the parameters $\hat{\lambda}$, $\hat{\vartheta}$, $\hat{\sigma}_o^2$, and $\hat{\sigma}_d^2$ were fixed to the values of the true underlying random walk parameters (see Methods section of the main article). The parameters $\hat{\mu}_1^{pre}$ and $\hat{\sigma}_1^{pre}$, specifying the mean and standard deviation of subjects' prior reward expectation for each bandit in the first trial, were fixed to $\hat{\mu}_1^{pre}$=50 and $\hat{\sigma}_1^{pre}$=4. Note that while these latter parameter values, which reflect reward expectancies on trial one, were chosen somewhat arbitrarily, they only influence modeled choice behavior on the first few trials and thus have low impact on the overall model fit (see Daw 2006).
